# SARS-CoV-2 Point Mutation and Deletion Spectra, and Their Association with Different Disease Outcome

**DOI:** 10.1101/2022.01.10.475768

**Authors:** Brenda Martínez-González, María Eugenia Soria, Lucía Vázquez-Sirvent, Cristina Ferrer-Orta, Rebeca Lobo-Vega, Pablo Mínguez, Lorena de la Fuente, Carlos Llorens, Beatriz Soriano, Ricardo Ramos, Marta Cortón, Rosario López-Rodríguez, Carlos García-Crespo, Isabel Gallego, Ana Isabel de Ávila, Jordi Gómez, Luis Enjuanes, Llanos Salar-Vidal, Jaime Esteban, Ricardo Fernandez-Roblas, Ignacio Gadea, Carmen Ayuso, Javier Ruíz-Hornillos, Nuria Verdaguer, Esteban Domingo, Celia Perales

**Affiliations:** Department of Clinical Microbiology, Instituto de Investigación Sanitaria-Fundación Jiménez Díaz University Hospital, Universidad Autónoma de Madrid (IIS-FJD, UAM) Av. Reyes Católicos 2, 28040 Madrid, Spain; Centro de Biología Molecular “Severo Ochoa” (CSIC-UAM), Consejo Superior de Investigaciones Científicas (CSIC), Campus de Cantoblanco, 28049 Madrid, Spain; Structural Biology Department, Institut de Biología Molecular de Barcelona CSIC, 08028 Barcelona, Spain; Department of Genetics & Genomics, Instituto de Investigación Sanitaria-Fundación Jiménez Díaz University Hospital, Universidad Autónoma de Madrid (IIS-FJD, UAM), Av. Reyes Católicos 2, 28040 Madrid, Spain; Centre for Biomedical Network Research on Rare Diseases (CIBERER), Instituto de Salud Carlos III, 28029 Madrid, Spain; Bioinformatics Unit, Instituto de Investigación Sanitaria-Fundación Jiménez Díaz University Hospital, Universidad Autónoma de Madrid (IIS-FJD, UAM), Madrid 28040, Spain; Biotechvana, “Scientific Park”, Universidad de Valencia, 46980 Valencia, Spain; Unidad de Genómica, “Scientific Park of Madrid”, Campus de Cantoblanco, 28049 Madrid, Spain; Centro de Investigación Biomédica en Red de Enfermedades Hepáticas y Digestivas (CIBERehd), Instituto de Salud Carlos III, 28029 Madrid, Spain; Instituto de Parasitología y Biomedicina ‘López-Neyra’ (CSIC), Parque Tecnológico Ciencias de la Salud, Armilla, 18016 Granada, Spain; Department of Molecular and Cell Biology, Centro Nacional de Biotecnología (CNB-CSIC), Consejo Superior de Investigaciones Científicas (CSIC), Campus de Cantoblanco, 28049 Madrid, Spain; Allergy Unit, Hospital Infanta Elena, Valdemoro, Madrid, Spain; Instituto de Investigación Sanitaria-Fundación Jiménez Díaz University Hospital, Universidad Autónoma de Madrid (IIS-FJD, UAM), Av. Reyes Católicos 2, 28040 Madrid, Spain; Faculty of Medicine, Universidad Francisco de Vitoria, Madrid, Spain

**Author notes:** Corresponding authors: Celia Perales and Esteban Domingo.

**Keywords:** COVID-19 severity, Mutant spectrum, Diversity index, Mutation, Deletion, nsp12 (polymerase), spike, Ultra-deep sequencing.

## Abstract

Mutant spectra of RNA viruses are important to understand viral pathogenesis, and response to selective pressures. There is a need to characterize the complexity of mutant spectra in coronaviruses sampled from infected patients. In particular, the possible relationship between SARS-CoV-2 mutant spectrum complexity and disease associations has not been established. In the present study, we report an ultra-deep sequencing (UDS) analysis of the mutant spectrum of amplicons from the nsp12 (polymerase)- and spike (S)-coding regions of thirty nasopharyngeal isolates (diagnostic samples) of SARS-CoV-2 of the first COVID-19 pandemic wave (Madrid, Spain, April 2020) classified according to the severity of ensuing COVID-19. Low frequency mutations and deletions, counted relative to the consensus sequence of the corresponding isolate, were overwhelmingly abundant. We show that the average number of different point mutations, mutations per haplotype and several diversity indices was significantly higher in SARS-CoV-2 isolated from patients who developed mild disease than in those associated with moderate or severe disease (exitus). No such bias was observed with RNA deletions. Location of amino acid substitutions in the three dimensional structures of nsp12 (polymerase) and S suggest significant structural or functional effects. Thus, patients who develop mild symptoms may be a richer source of genetic variants of SARS-CoV-2 than patients with moderate or severe COVID-19.

**IMPORTANCE:** The study shows that mutant spectra of SARS-CoV-2 from diagnostic samples differ in point mutation abundance and complexity, and that significantly larger values were observed in virus from patients who developed mild COVID-19 symptoms. Mutant spectrum complexity is not a uniform trait among isolates. The nature and location of low frequency amino acid substitutions present in mutant spectra anticipate great potential for phenotypic diversification of SARS-CoV-2.

## INTRODUCTION

Betacoronavirus SARS-CoV-2 emerged in the human population in 2019, and it is the causal agent of the new pandemic disease COVID-19 (1), with a death toll which is increasing at the time of this writing (https://covid19.who.int/). Genetic variations in SARS-CoV-2 genomes [annotated in the GISAID (https://www.gisaid.org/), PubMed (https://www.ncbi.nlm.nih.gov/pmc/), and ENA data banks (https://www.ebi.ac.uk/ena/browser/home); among others] affect non-structural and structural protein-coding regions. Despite the short history of SARS-CoV-2 circulation, newly arising variants exhibiting different mutational patterns are regularly being identified. A distinction has been made between variants of interest (VOI), due to features with potential impact (such as transmissibility), and variants of concern (VOC), due to definite evidence of enhanced transmissibility (https://www.who.int/en/activities/tracking-SARS-CoV-2-variants/). New SARS-CoV-2 variants are likely to become prominent as COVID-19 continues, despite natural or vaccine-induced immunity (2–5). Likewise, the generation of viral escape mutants is a major concern as a potential limitation of immune and antiviral agent efficacy for SARS-CoV-2 (6–10), as it has been established for other RNA viruses.

The first step in the diversification of viruses during their epidemic spread is the generation of variants within each infected host. This pattern of intra-host evolution results in the formation of mutant spectra that constitute reservoirs of genetic and phenotypic virus variants in the infected host (11, 12). Studies with several RNA viruses have shown that viral intra-mutant spectrum complexity, estimated by the average number of mutations per genome, expressed by a series of diversity indices [Shannon entropy, maximum mutation frequency, Gini Simpson, nucleotide diversity, number of polymorphic sites, and number of haplotypes (13, 14)] may have an impact on viral tropism, viral persistence, disease progression and response to antiviral interventions [several cases have been described or reviewed in (11, 15–22)]. Evidence of quasispecies dynamics has been reported for SARS-CoV-2 (23–29), as well as for other coronaviruses (30–34). However, it is unclear how mutant spectrum complexity parameters of this emerging pathogen vary among different viral isolates, and whether previously observed effects of mutant spectrum composition on RNA virus behavior apply also to SARS-CoV-2, particularly its connection with disease severity.

Two recent studies indicated higher mutant spectrum complexity in SARS-CoV-2 from patients who developed severe disease than mild disease, either analyzing the spike (S)-coding regions (35), or the entire genome with limited mutant spectrum resolution (36). In the present study, we have examined mutant spectra of the nsp12 (polymerase)- and S-coding regions of SARS-CoV-2 present in 30 nasopharyngeal swab samples taken at the time of diagnosis of patients progressing towards disparate disease outcomes. Applying a 0.5% cut-off value for point mutation and deletion detection, using SeekDeep as bioinformatics platform, we found that virus from patients who developed mild disease exhibited a significantly higher mutant spectrum complexity than virus from patients who developed moderate or severe disease (exitus). The difference occurred both in the nsp12 (polymerase)- and S-coding regions. In contrast, no significant differences in the spectrum of minority deletions were observed among virus from the three patient’s categories (mild, moderate or severe disease). Some amino acid substitutions found at low frequency in mutant spectra, including substitutions with low statistical acceptability and with potential functional effects, are nevertheless present in SARS-CoV-2 isolates recorded in data banks.

## RESULTS

### SARS-CoV-2 mutant spectra from patients progressing towards different COVID-19 severity

We previously classified 448 patients [Fundación Jiménez Díaz (FJD) cohort, Madrid, Spain, April 2020] according to the COVID-19 severity into three categories: mild, moderate and severe COVID-19 ─based on a number of demographic and clinical parameters─ and we found a positive association between viral load in nasopharyngeal swabs and disease severity (37). For the present study, we have chosen thirty of the nasopharyngeal samples based on three criteria: (i) the COVID-19 category, including 10 patients who developed mild symptoms, 10 patients who developed moderate disease, and 10 patients who progressed to severe disease and exitus; (ii) patients whose diagnostic (RT-PCR RNA samples) displayed similar Ct values (average Ct=25.37±3.9 for mild, Ct=21.81±2.4 for moderate, and Ct=20.38±2.9 for exitus patients); and (iii) similar time interval between symptom onset and swab collection (average 5.78±4.2 days for mild, 4.89±3.1 days for moderate and 4.5±2.6 days for exitus patients). When present, comorbidities were equally represented among the different COVID-19 severities (Table S1 in https://saco.csic.es/index.php/s/8GH5aJgritCjEx5).

To set up ultra-deep sequencing (UDS) analyses of SARS-CoV-2 obtained from nasopharyngeal swabs, we have adapted experimental protocols previously used for HCV quasispecies characterization (38–41), and applied the SeekDeep pipeline (42) to the analysis of minority point mutations and deletions in SARS-CoV-2 mutant spectra (described in Materials and Methods). RNA from nasopharyngeal swabs was extracted, amplified and subjected to UDS using MiSeq platform (Illumina). Four amplicons (A1 to A4) covering nucleotides 14,534 to 16,054 of the nsp12 (polymerase)-coding region, and two amplicons (A5 and A6) covering nucleotides 22,872 to 23,645 of the S-coding region were analyzed (Fig. 1). The total number of clean reads was 19,592,197, corresponding to 653,073 (range 316,710-910,727) reads per patient, that yielded an average of 110,689 (range 38,865-215,662) clean reads per amplicon, with a 0.5% cut-off frequency for point mutations and deletions (Fig. S1 in https://saco.csic.es/index.php/s/8GH5aJgritCjEx5).

**FIG 1.**
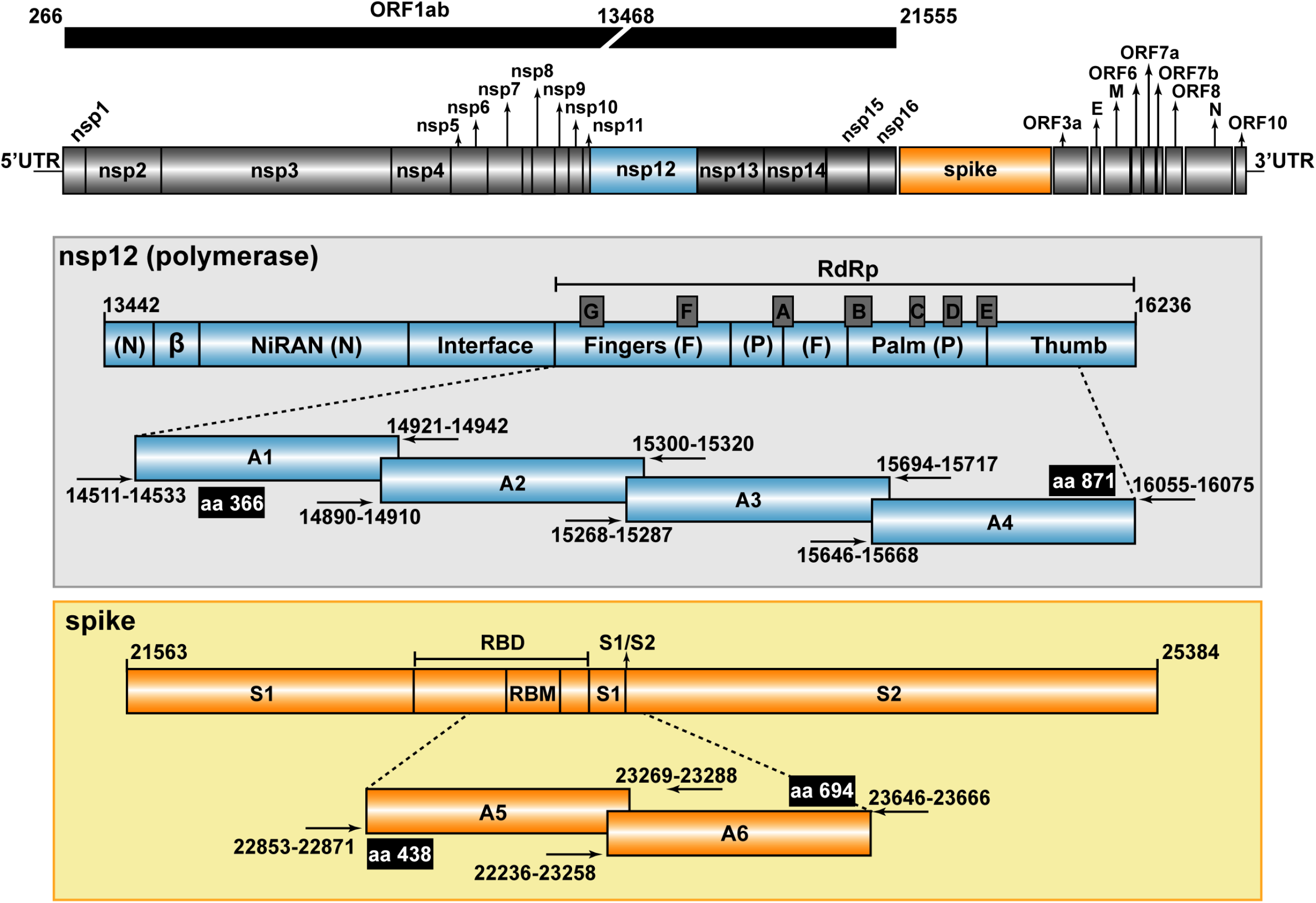
Representation of the SARS-CoV-2 genome, encoded proteins, and amplicons analyzed by UDS. The region corresponding to the ORF1ab of the virus is shown at the top. In the two boxes, the nsp12 (blue) and spike (orange) have been expanded, with the first and last nucleotide number given at the beginning and the end of the bars, respectively (genome numbering is according to the reference genome NCBI accession number: NC_045512.2). Relevant protein domains are indicated, including motifs A to G depicted as protruding grey boxes in nsp12 (polymerase), and the receptor binding motif (RBM) and the S1/S2 cleavage site in S. The amplicons [A1 to A4 for the nsp12 (polymerase) and A5, A6 for S] are shown flanked by horizontal arrows that mark the position of the oligonucleotide primers used for amplification (oligonucleotide sequences are given in Table S5 in https://saco.csic.es/index.php/s/8GH5aJgritCjEx5). Flanking black boxes indicate the amino acids (aa) of nsp12 (polymerase) and S covered by the amplicons.

To provide a general picture of SARS-CoV-2 divergence and mutant spectrum heterogeneity, we constructed a heat map representing the frequency of each variation in the nsp12 (polymerase) and S-coding regions (point mutations and deletions; no insertions were detected), relative to the genomic sequence of a Wuhan isolate (identified as NCBI reference sequence NC_045512.2), and divided the samples according to different COVID-19 severity (Fig. 2 and Table S2 in https://saco.csic.es/index.php/s/8GH5aJgritCjEx5). Considering all patients analyzed, the number of positions that included a variation (either a point mutation or a deletion) was two-fold higher in the S-coding region (105 positions with a genomic modification out of 774 positions analyzed) than in the nsp12 (polymerase)-coding region (91 positions modified out of 1,521 positions analyzed). In addition to minority mutations in each mutant spectrum, a total of six different dominant mutations relative to the reference sequence (those with frequencies between 90% and 100%) were also present; they are identified as “Divergence” in Fig. 2. This class of mutations has been excluded for the quantification of mutations and complexity indices in a mutant spectrum. Ninety-four percent of mutations were found at frequencies that ranged between 0.5% and 30% within its mutant spectrum, whereas only 6% corresponded to “Divergence” mutations (p<0.001; proportion test). Interestingly, 62 out of 97 point mutations (64%) within the mutant spectra were detected at frequencies below 2% (Fig. 2).

**FIG 2.**
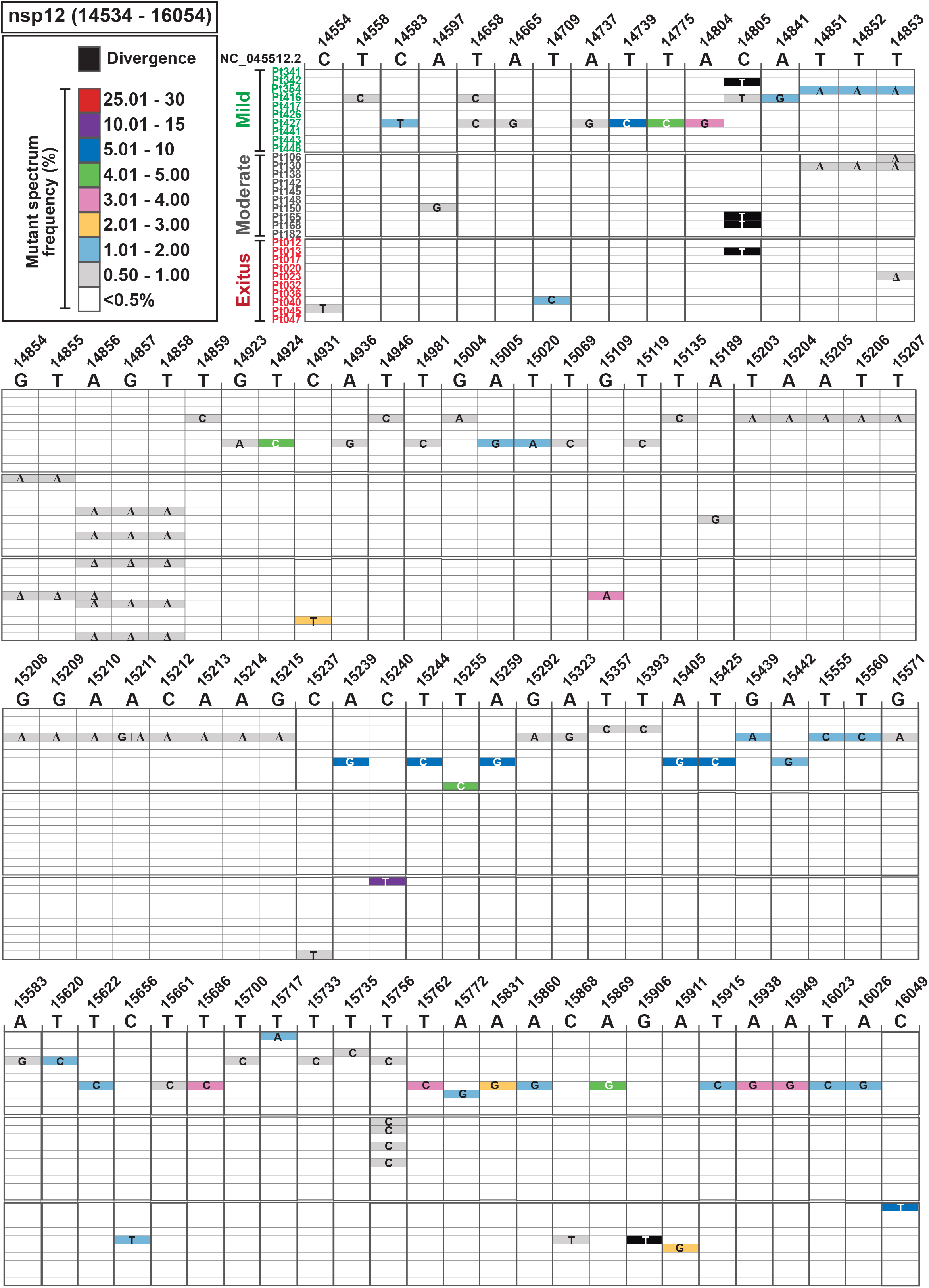
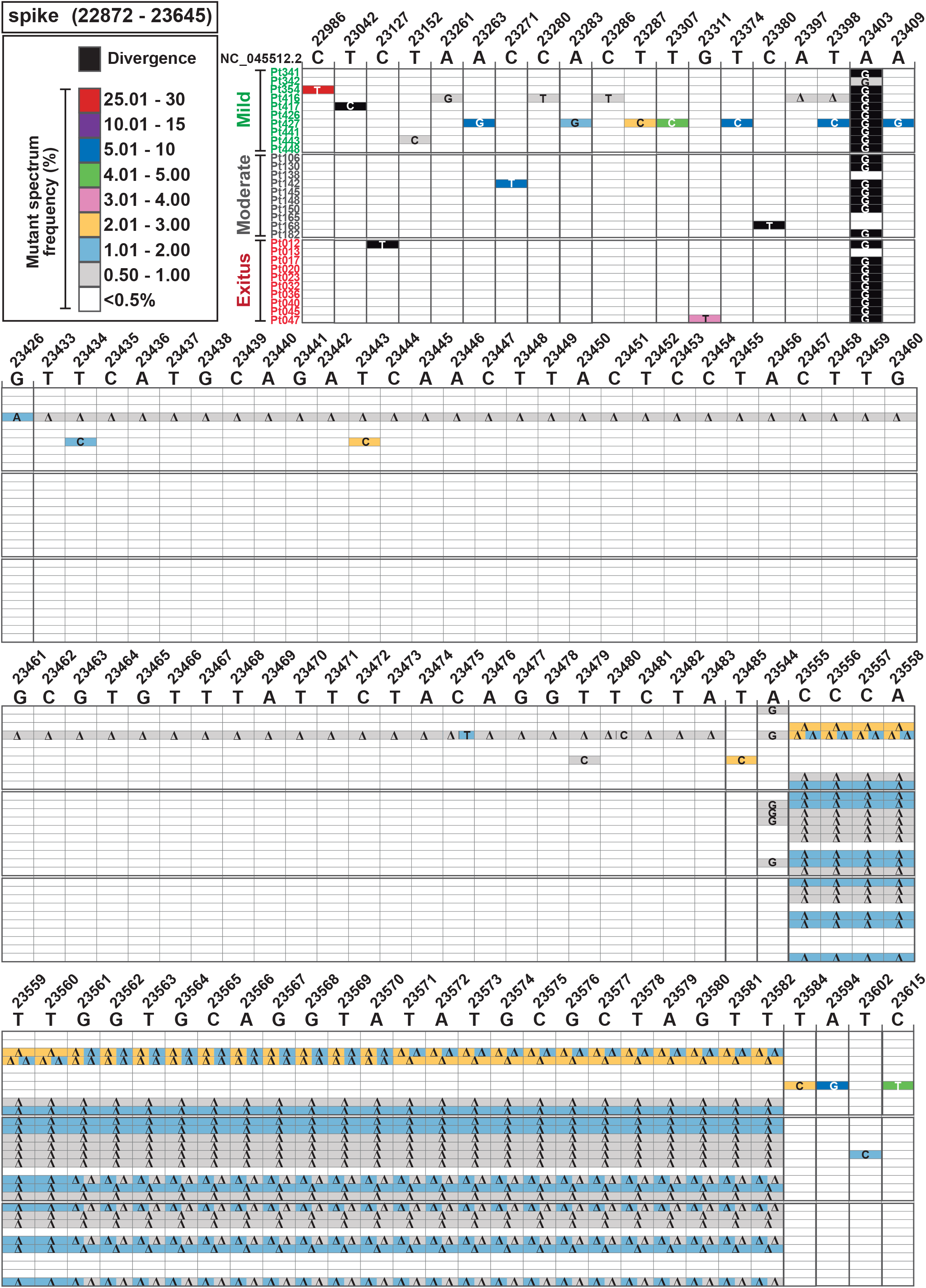
Heat map of point mutation and deletion frequencies in mutant spectra of SARS-CoV-2 from individual patients. Data are presented in two blocks, one for the nsp12 (polymerase)-coding region (genomic residues 14,534-16,054), and another for the S-coding region (genomic residues 22,872-23,645). Only positions with a mutation or those affected by a deletion are represented. Each row corresponds to a patient, and patients have been divided in those with mild, moderate and exitus disease outcomes (color coded, and with the patient identification code written at the left of each row). The patients’ clinical status and demographic data are described in Table S1 in https://saco.csic.es/index.php/s/8GH5aJgritCjEx5. Mutations and deletions have been identified relative to NCBI reference sequence NC_045512.2. Each mutation and deletion (delta symbol Δ) with a frequency above the cut-off level (0.5%) is indicated, and its frequency within the mutant spectrum retrieved from each patient has been visualized with a color code displayed in the heading boxes (top left of the two blocs). Procedures are detailed in Materials and Methods.

To evaluate if some parameters of the mutant spectra (considering only point mutations present at a frequency below 30%) were associated with COVID-19 severity, we first counted the number of different point mutations present in virus from each patient group. In the two coding regions analyzed, the average number of different mutations in virus from patients with mild disease was significantly higher than in virus from patients with moderate disease or exitus [p < 0.001 for the comparison between mild versus moderate and mild versus exitus, both for nsp12 (polymerase)- and S-coding region; proportion test]; no significant difference was noted between moderate and exitus patients [p = 0.081 and p = 0.603 for nsp12 (polymerase)- and S-coding regions, respectively; proportion test]; normalization of the number of different mutations to the length of the regions analyzed did not modify the result (Fig. 3A). No such difference among patient groups was observed with the number of different deletions (all p-values > 0.05; proportion test), although a trend towards a larger number of deletions in virus from patients who developed mild disease was maintained in the S-coding region (Fig. 3B). Thus, SARS-CoV-2 mutant spectra from diagnostic samples of patients who evolved to mild disease included a significantly larger average number of mutations, but not of deletions, than virus from patients who progressed towards moderate or severe (exitus) COVID-19.

**FIG 3.**
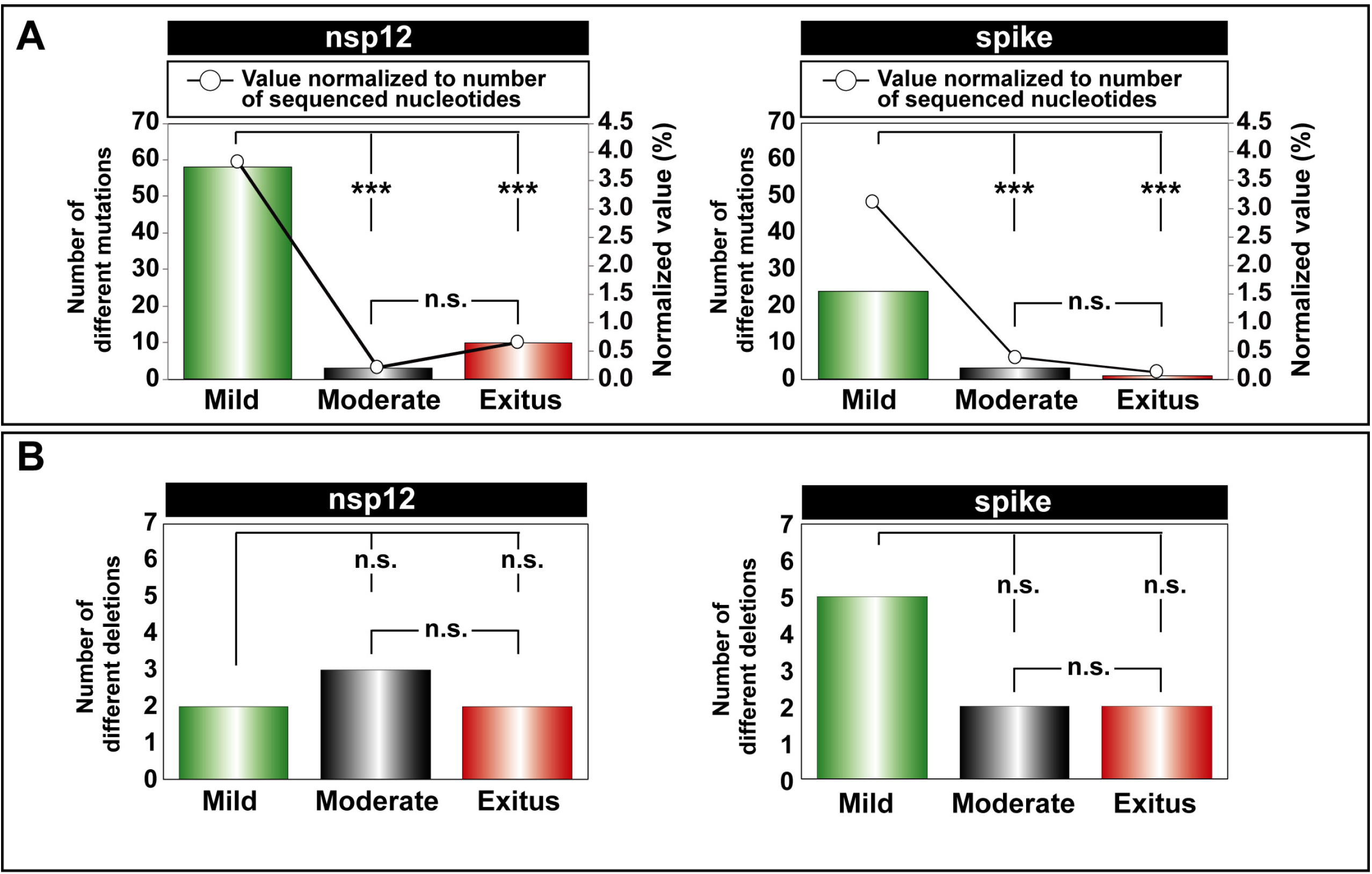
Point mutations and deletions in the mutant spectra of SARS-CoV-2 isolates, distributed according to COVID-19 severity. The point mutations and deletions are those depicted in Fig. 2. **(A)** Total number of different point mutations in the nsp12 (polymerase)- (left panel) and the S- (right panel) coding region distributed according to disease severity (mild, moderate, exitus, as indicated in the abscissa) in the patients from whom the virus was isolated. Bars indicate the total absolute number of mutations (left ordinate axes) and empty dots give the percentages normalized to the length in nucleotides of the sequenced regions (right ordinate axes). **(B)** Total number of different deletions in the nsp12 (polymerase)- (left panel) and the S- (right panel) coding region distributed according to disease severity in the patients from whom the virus was isolated. For (A) and (B) the statistical significance of the differences was determined by the proportion test; ns; not significant, ***p<0.001.

### Evaluation of complexity indices

The comparison of SARS-CoV-2 mutant spectra was extended to two groups of diversity indices: abundance (which consider the reads of entities and their frequency in the mutant spectrum), and incidence (which consider only reads of entities) (13). To this aim, we have adapted the QSutils package (43) to the quantification of diversity indices for SARS-CoV-2 mutant spectra (described in Materials and Methods). In the nsp12 (polymerase)-coding region, a significant increase of the values of abundance and incidence indices was observed in samples from patients who developed mild disease, as compared with samples from patients with moderate disease (p < 0.001 for H_S_, H_GS_, Mf_max_ and π; p=0.001 for number of polymorphic sites and number of haplotypes; Wilcoxon test). Also significant was the difference between samples associated with mild disease and severe disease (exitus) (p = 0.004 for H_S_, p = 0.010 for H_GS_, p = 0.012 for Mf_max_ and p = 0.010 for π; p = 0.004 for number of polymorphic sites and number of haplotypes; Wilcoxon test). The same tendency was observed in the S-coding region but the differences did not reach statistical significance (all p-values > 0.05; proportion test) (Fig. 4 and Table S3 in https://saco.csic.es/index.php/s/8GH5aJgritCjEx5). In each amplicon, a larger number of haplotypes was found in samples associated with mild than moderate or severe disease, and the majority of mutated haplotypes included only one mutation (Fig. S2 in https://saco.csic.es/index.php/s/8GH5aJgritCjEx5). Thus, the higher abundance of mutations in SARS-CoV-2 mutant spectra from patients who exhibited only mild symptoms is also reflected in an increase of mutant spectrum complexity.

**FIG 4.**
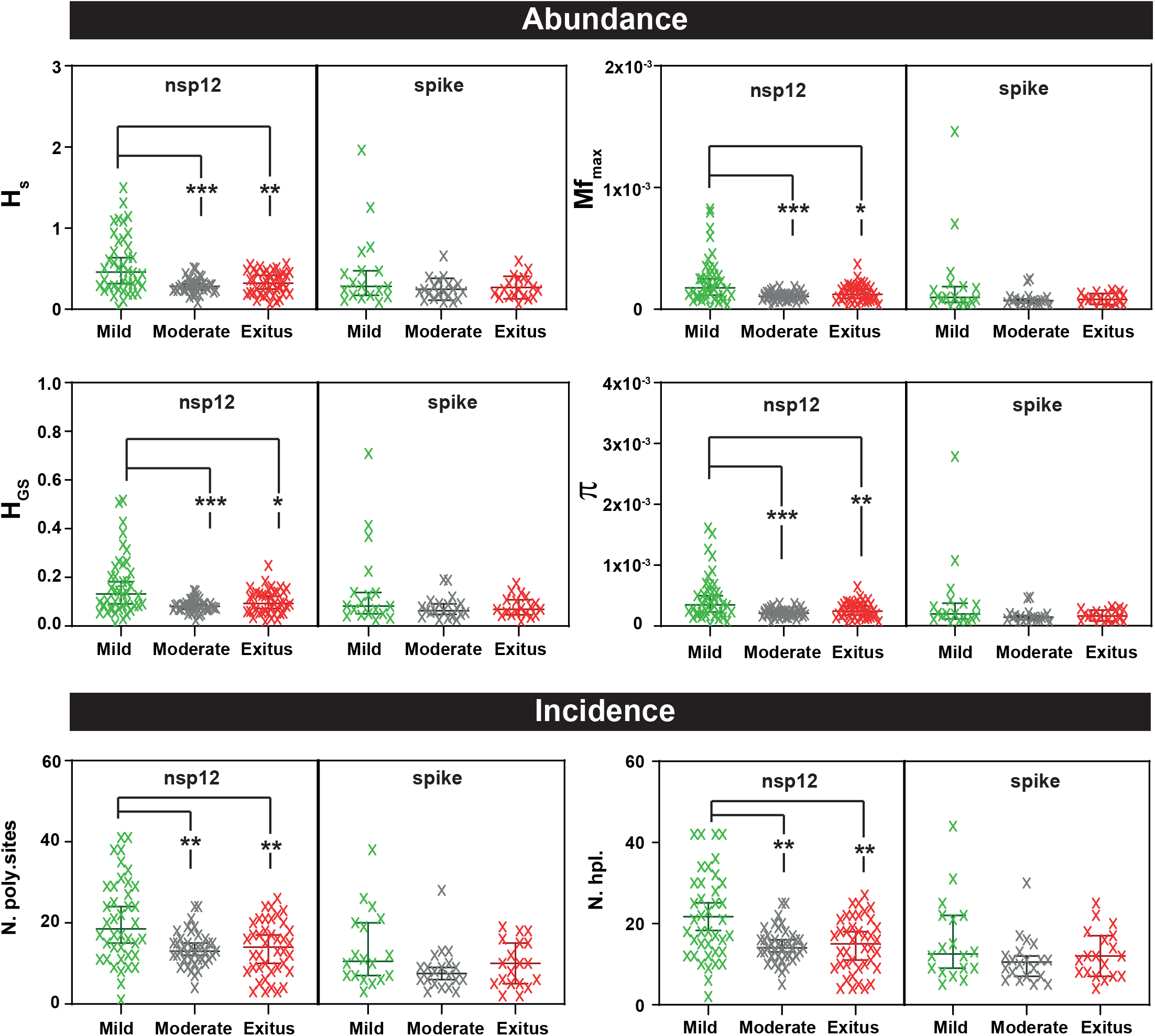
Comparison of the diversity indices for all amplicons of either the nsp12 (polymerase)- or S-coding region, distributed according to virus-associated disease severity. The types of indices (abundance or incidence) are indicated in the heading filled boxes. The specific index is indicated in ordinate (13) (H_s_, Shannon entropy; Mf_max_, maximum mutation frequency; H_GS_, Gini Simpson; π, nucleotide diversity; N. poly.sites, number of polymorphic sites; N. hpl., number of haplotypes). Each cross is the numerical value obtained for the virus of an individual patient; patients have been distributed according to disease severity as indicated in abscissa (color coded). Data were obtained using a cut-off value of 0.1%, as previously reported (13). Values for each amplicon and patient are compiled in Table S3 in https://saco.csic.es/index.php/s/8GH5aJgritCjEx5. The statistical significance of the differences has been determined by the Wilcoxon test. *, p<0.05; **, p<0.01; ***p<0.001; absence of connecting lines means that the difference between two patient groups was not statistically significant.

### Point mutation and amino acid substitution types in SARS-CoV-2 mutant spectra

Considering mutant spectra of all samples analyzed, transitions and non-synonymous mutations were more abundant than transversions and synonymous mutations, respectively, with different degrees of statistical significance (Table 1); a similar trend was also observed when the samples were divided according to COVID-19 severity of the patients.

**Table 1.** Point mutations in the mutant spectra of SARS-CoV-2 isolates^a^.

|  |  | **Disease severity** |  |  |  |
| --- | --- | --- | --- | --- | --- |
|  |  | **Total** | **Mild** | **Moderate** | **Exitus** |
| **nsp12** | **Transitions (%)** | 68 (97.14%) | 56 (96.55%) | 3 (100%) | 10 (100%) |
|  | **Transversions (%)** | 2 (2.86%) | 2 (3.45%) | 0 (0%) | 0 (0%) |
|  | **p-value** | <0.001 | <0.001 | 0.051 | <0.001 |
|  | **Significance<sup>b</sup>** | *** | *** | n.s. | *** |
|  | **Synonymous (%)** | 29 (41.43%) | 24 (41.38%) | 2 (66.67%) | 4 (40%) |
|  | **Non-Synonymous (%)** | 41 (58.57%) | 34 (58.62%) | 1 (33.33%) | 6 (60%) |
|  | **p-value** | 0.031 | 0.047 | 0.5 | 0.327 |
|  | **Significance<sup>b</sup>** | * | * | n.s. | n.s. |
| **spike** | **Transitions (%)** | 26 (96.30%) | 24 (100%) | 3 (100%) | 0 (0%) |
|  | **Transversions (%)** | 1 (3.70%) | 0 (0%) | 0 (0%) | 1 (100%) |
|  | **p-value** | <0.001 | <0.001 | 0.051 | 0.5 |
|  | **Significance<sup>b</sup>** | *** | *** | n.s. | n.s. |
|  | **Synonymous (%)** | 11 (40.74%) | 10 (41.67%) | 1 (33.33%) | 0 (0%) |
|  | **Non-Synonymous (%)** | 16 (59.26%) | 14 (58.33%) | 2 (66.67%) | 1 (100%) |
|  | **p-value** | 0.138 | 0.193 | 0.5 | 0.051 |
|  | **Significance<sup>b</sup>** | n.s. | n.s. | n.s. | n.s. |

<sup>a</sup> Different number of point mutations distributed according to COVID-19 severity in the nsp12 (polymerase)- and spike-coding region.
<sup>b</sup> The statistical significance of the differences (n.s., not significant; \* p<0.05; \*\*\* p<0.001) was calculated using the proportion test.

In the nsp12 (polymerase)-coding region, the frequency of mutation types normalized to base composition ranked as follows: T to C > A to G > C to T; when dividing the samples according to disease severity, the most frequent mutation in exitus patients was C to T (Fig. S3A in https://saco.csic.es/index.php/s/8GH5aJgritCjEx5). In the S-coding region the ranking was T to C > A to G = C to T (Fig. S3B in https://saco.csic.es/index.php/s/8GH5aJgritCjEx5). T to C transitions were the most frequent mutation type in the third codon position (67.50%), whereas A to G was the most prevalent type at the second and first codon positions (45.16% and 38.46%, respectively).

The amino acid substitutions found in nsp12 (polymerase) and S were positioned in the three-dimensional structure of the proteins [Protein Data Bank (http://www.wwpdb.org/)], their statistical acceptability was evaluated with PAM250 matrix (44), and their potential functional effects was estimated by applying the SNAP2 predictor (45). All amino acid substitutions found in nsp12 (polymerase) and S are listed in Table S2 in https://saco.csic.es/index.php/s/8GH5aJgritCjEx5, together with their PAM250 and SNAP2 scores; their location in the three dimensional structure of the proteins is depicted in Fig. S4 in https://saco.csic.es/index.php/s/8GH5aJgritCjEx5. Those amino acid substitutions which suggest alteration of protein structure or function are described in Tables 2 and 3. Some of the substitutions in nsp12 (polymerase) predict positive or negative functional effects (Table 2 and Fig. 5). For example, V557I may enhance the stability of the interaction with nitrogen base T+1, and Q822H predicts increased stability of loop in the thumb domain. In contrast, D618N abolishes the catalytic aspartate of polymerase in domain A, and C765R should distort the catalytic domain (Table 2 and Fig. 5). The amino acid substitutions observed in S tend to increase the hydrophobicity of the region where they are located (Table 3 and Fig. 6). The replacement of A by V at position 475 may enhance interactions of S with ACE2; A522V may contribute to stabilize the RBD domain in the “open” position through contacts with neighbor V, T, P and L residues; R567G could facilitate fusion with the host cell; A570V may bring closer two S chains (Table 3 and Fig. 6). Drastic substitutions may belong to defective genomes that have a transient existence or that may be maintained by complementation (see Discussion).

**Table 2.**
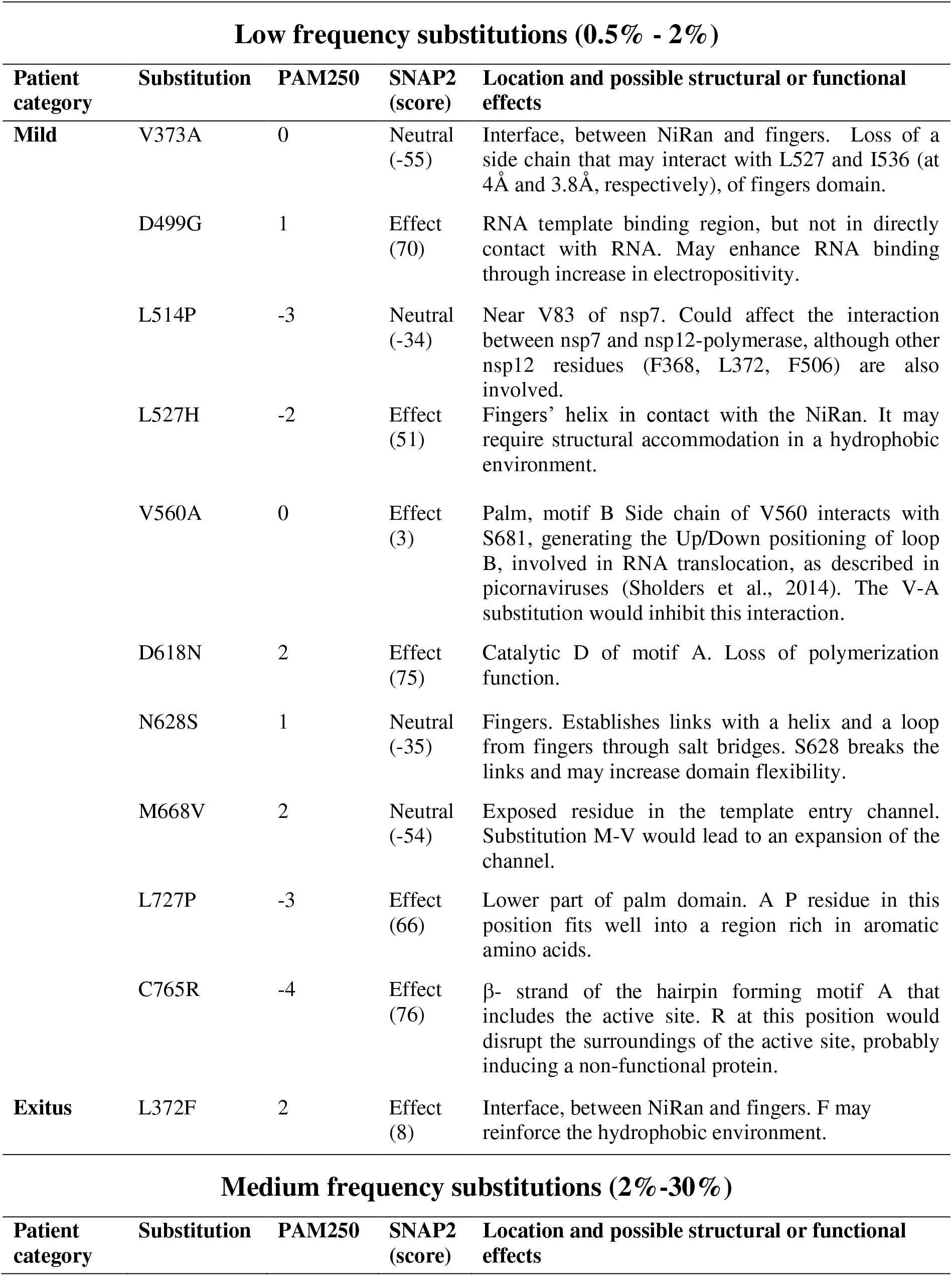
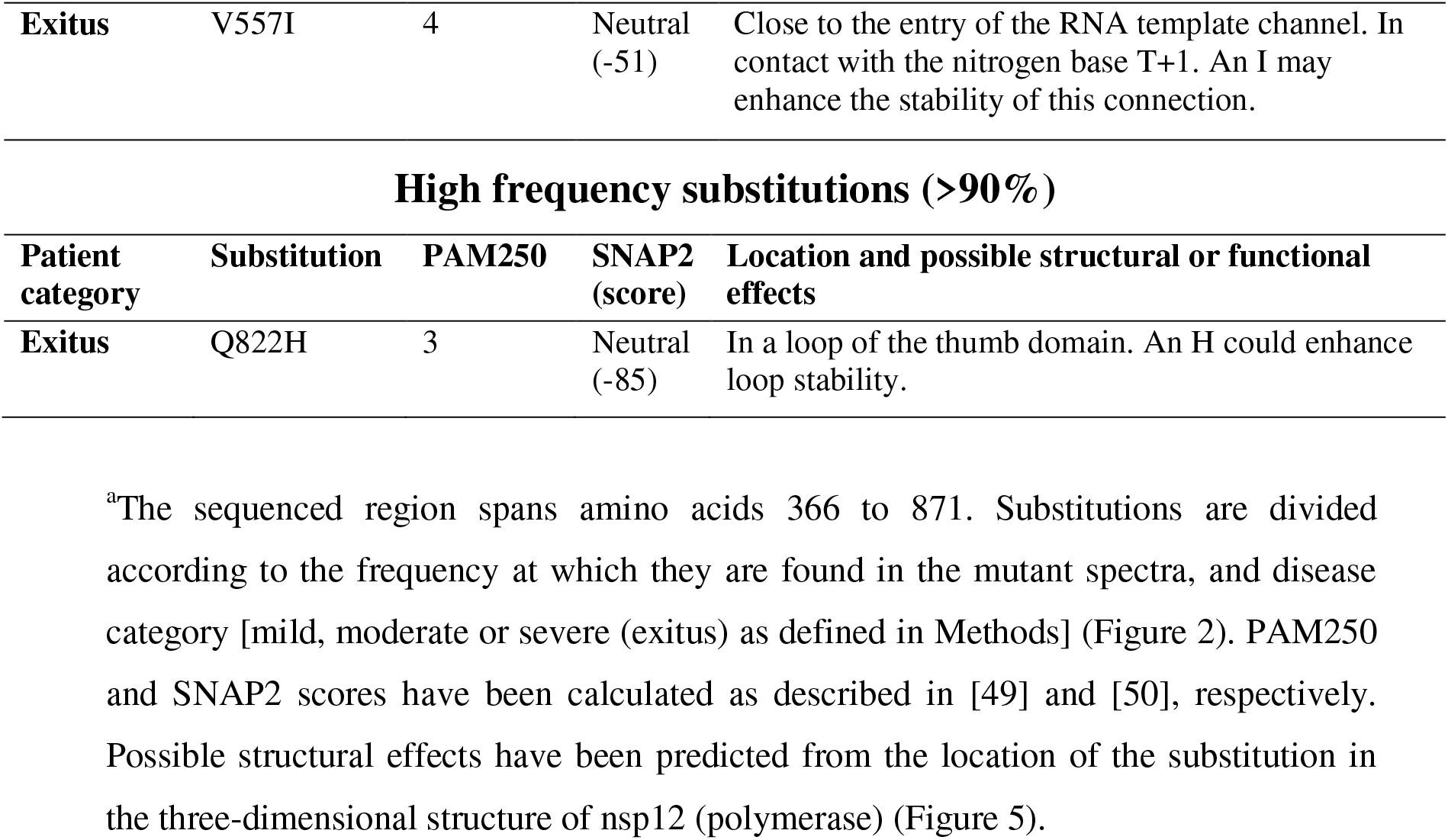
Amino acid substitutions at the nsp12 (polymerase) in the mutant spectra of SARS-CoV-2^a^.

**Table 3.**
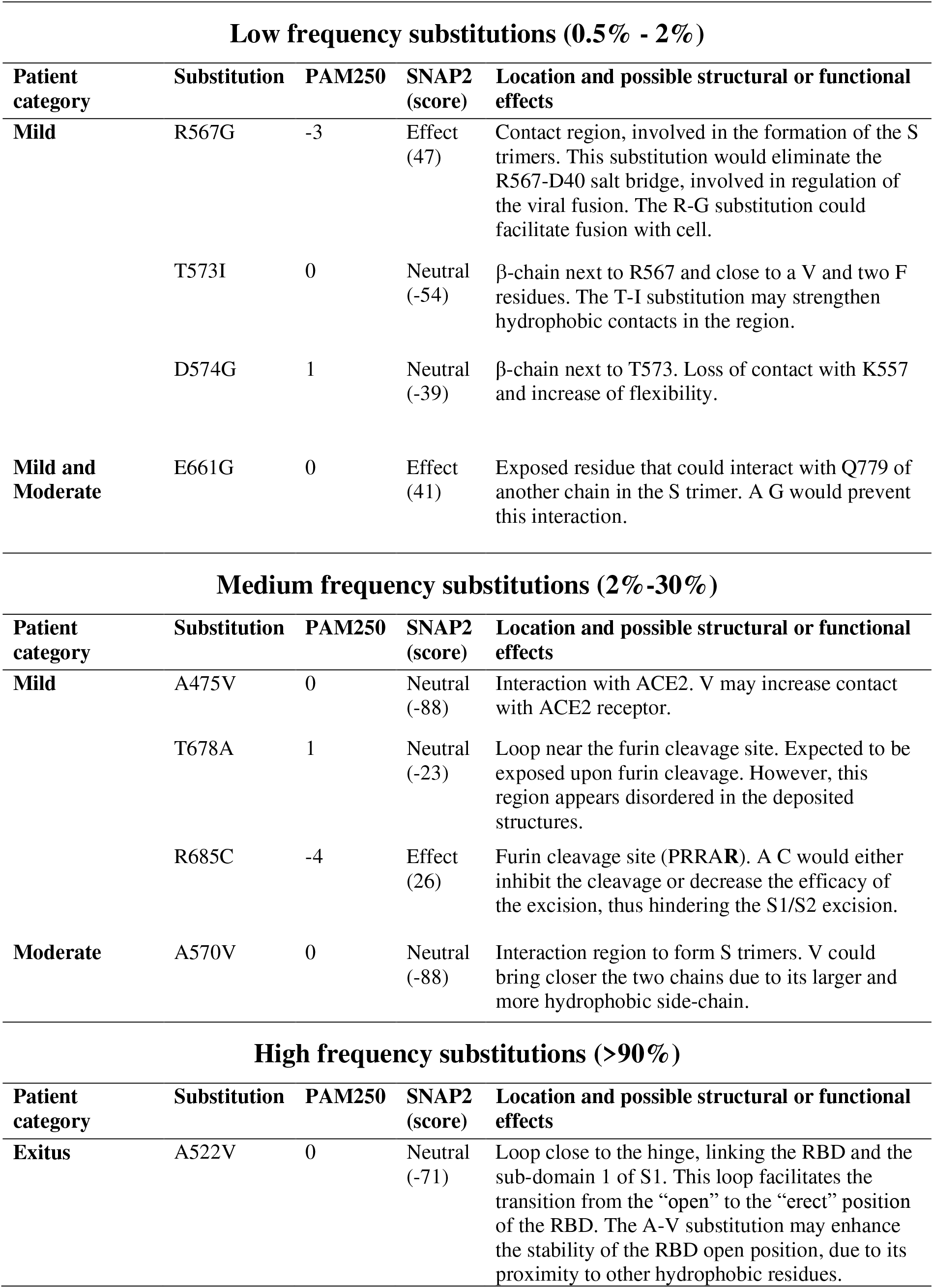
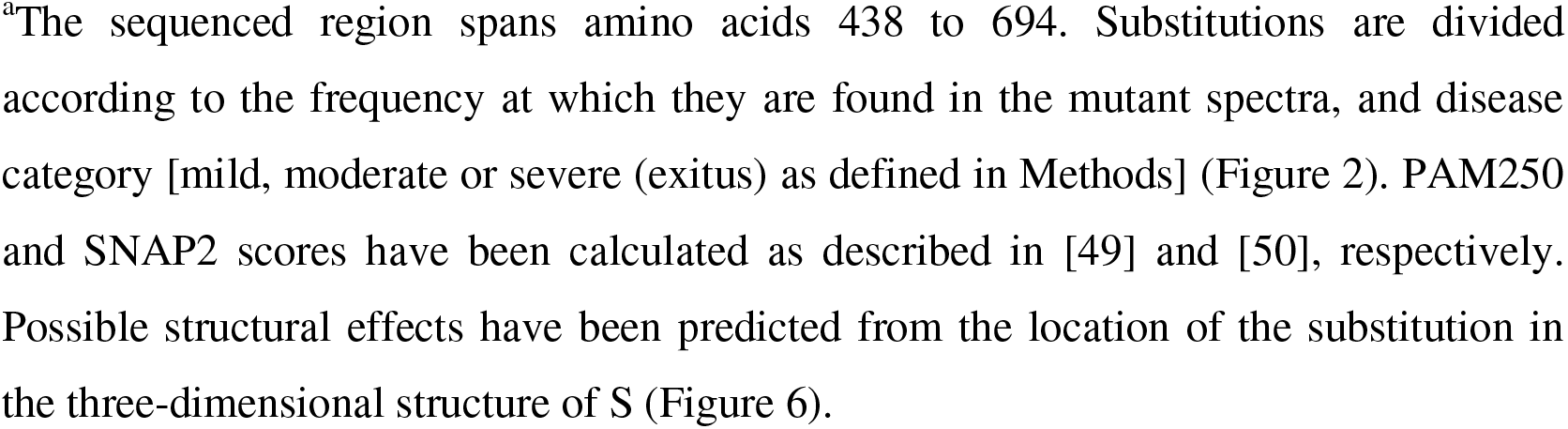
Amino acid substitutions at the spike (S) protein in the mutant spectra of SARS-CoV-2^a^.

| **Low frequency substitutions (0.5% - 2%)** |  |  |  |  |
| --- | --- | --- | --- | --- |
| **Patient category** | **Substitution** | **PAM250** | **SNAP2 (score)** | **Location and possible structural or functional effects** |
| **Mild** | R567G | -3 | Effect (47) | Contact region, involved in the formation of the S trimers. This substitution would eliminate the R567-D40 salt bridge, involved in regulation of the viral fusion. The R-G substitution could facilitate fusion with cell. |
|  | T573I | 0 | Neutral (-54) | β-chain next to R567 and close to a V and two F residues. The T-I substitution may strengthen hydrophobic contacts in the region. |
|  | D574G | 1 | Neutral (-39) | β-chain next to T573. Loss of contact with K557 and increase of flexibility. |
| **Mild and Moderate** | E661G | 0 | Effect (41) | Exposed residue that could interact with Q779 of another chain in the S trimer. A G would prevent this interaction. |
| **Medium frequency substitutions (2%-30%)** |  |  |  |  |
| **Patient category** | **Substitution** | **PAM250** | **SNAP2 (score)** | **Location and possible structural or functional effects** |
| **Mild** | A475V | 0 | Neutral (-88) | Interaction with ACE2. V may increase contact with ACE2 receptor. |
|  | T678A | 1 | Neutral (-23) | Loop near the furin cleavage site. Expected to be exposed upon furin cleavage. However, this region appears disordered in the deposited structures. |
|  | R685C | -4 | Effect (26) | Furin cleavage site (PRRAR). A C would either inhibit the cleavage or decrease the efficacy of the excision, thus hindering the S1/S2 excision. |
| **Moderate** | A570V | 0 | Neutral (-88) | Interaction region to form S trimers. V could bring closer the two chains due to its larger and more hydrophobic side-chain. |
| **High frequency substitutions (>90%)** |  |  |  |  |
| **Patient category** | **Substitution** | **PAM250** | **SNAP2 (score)** | **Location and possible structural or functional effects** |
| **Exitus** | A522V | 0 | Neutral (-71) | Loop close to the hinge, linking the RBD and the sub-domain 1 of S1. This loop facilitates the transition from the “open” to the “erect” position of the RBD. The A-V substitution may enhance the stability of the RBD open position, due to its proximity to other hydrophobic residues. |

<sup>a</sup>The sequenced region spans amino acids 438 to 694. Substitutions are divided according to the frequency at which they are found in the mutant spectra, and disease category [mild, moderate or severe (exitus) as defined in Methods] (Figure 2). PAM250 and SNAP2 scores have been calculated as described in [49] and [50], respectively. Possible structural effects have been predicted from the location of the substitution in the three-dimensional structure of S (Figure 6).

**FIG 5.**
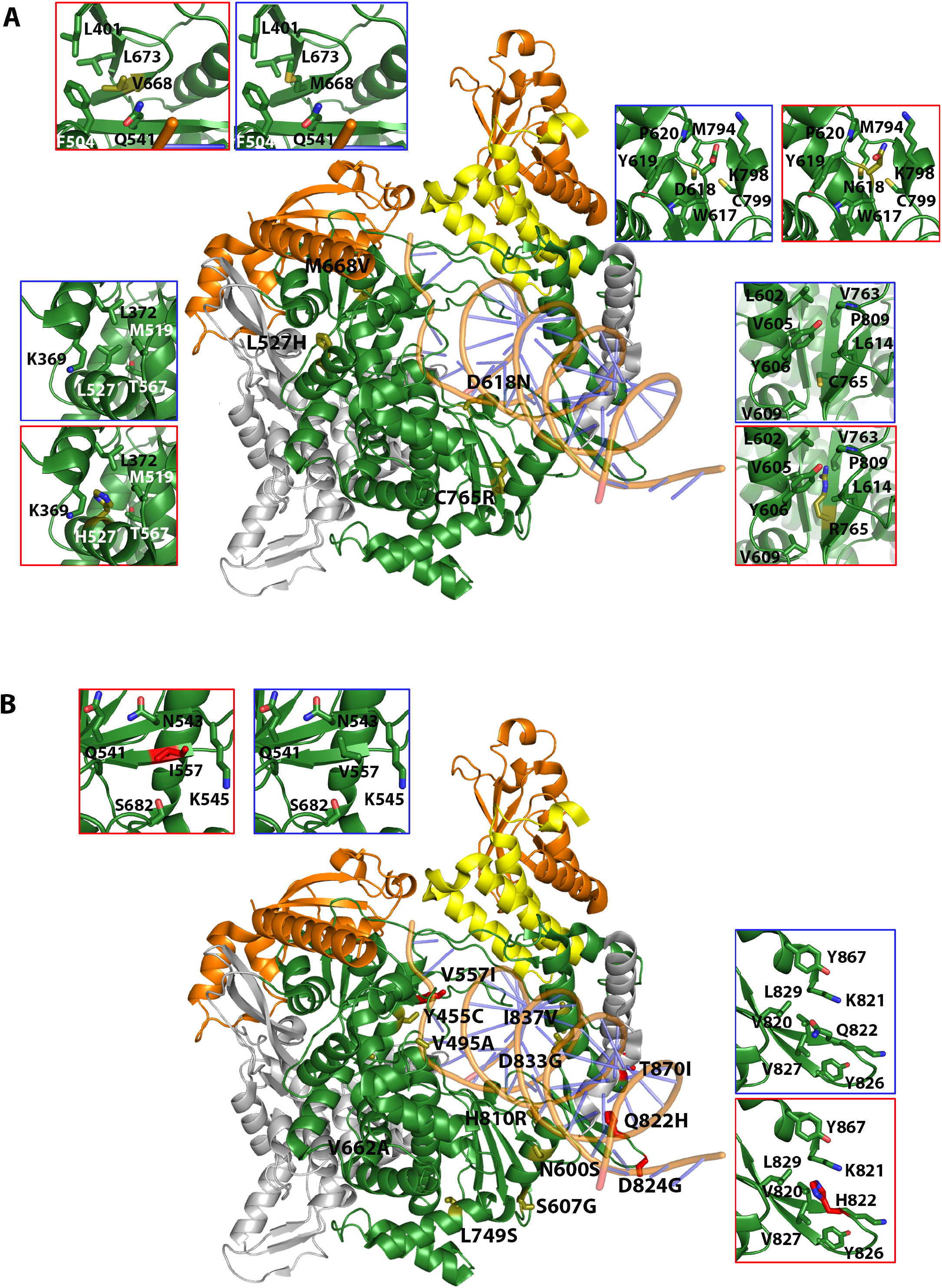
Location of amino acid substitutions in the three-dimensional structure of nsp12 (polymerase). The structure used as reference is that of the replication complex nsp12-nsp8-nsp7 (PDB code 6NUR with the RNA superimposed from 7CYQ). **(A)** Substitutions found at low frequency (0.5% to 2%) in the mutant spectra. The central structure is a cartoon representation of the nsp12, depicted in grey and green, the latter showing the regions covered by amplicons A1-A4 (indicated in Fig. 1). Contact proteins nsp8 (orange) and nsp7 (yellow) are also drawn. Substitutions are labeled, and amino acids are shown as sticks in different colors, according to associated disease category: exitus in red; mild disease in yellow. Insets highlight the interactions of some substitutions with neighboring residues within a 5-Å radius. Two insets are shown per position, indicating the original and mutated residues, squared in blue and red, respectively. (B) Same design as A but with substitutions found at frequency higher than 2%. The substitutions, their frequency in the mutant spectrum, acceptability, functional score, and possible structural or functional effects are listed in Table 2.

**FIG 6.**
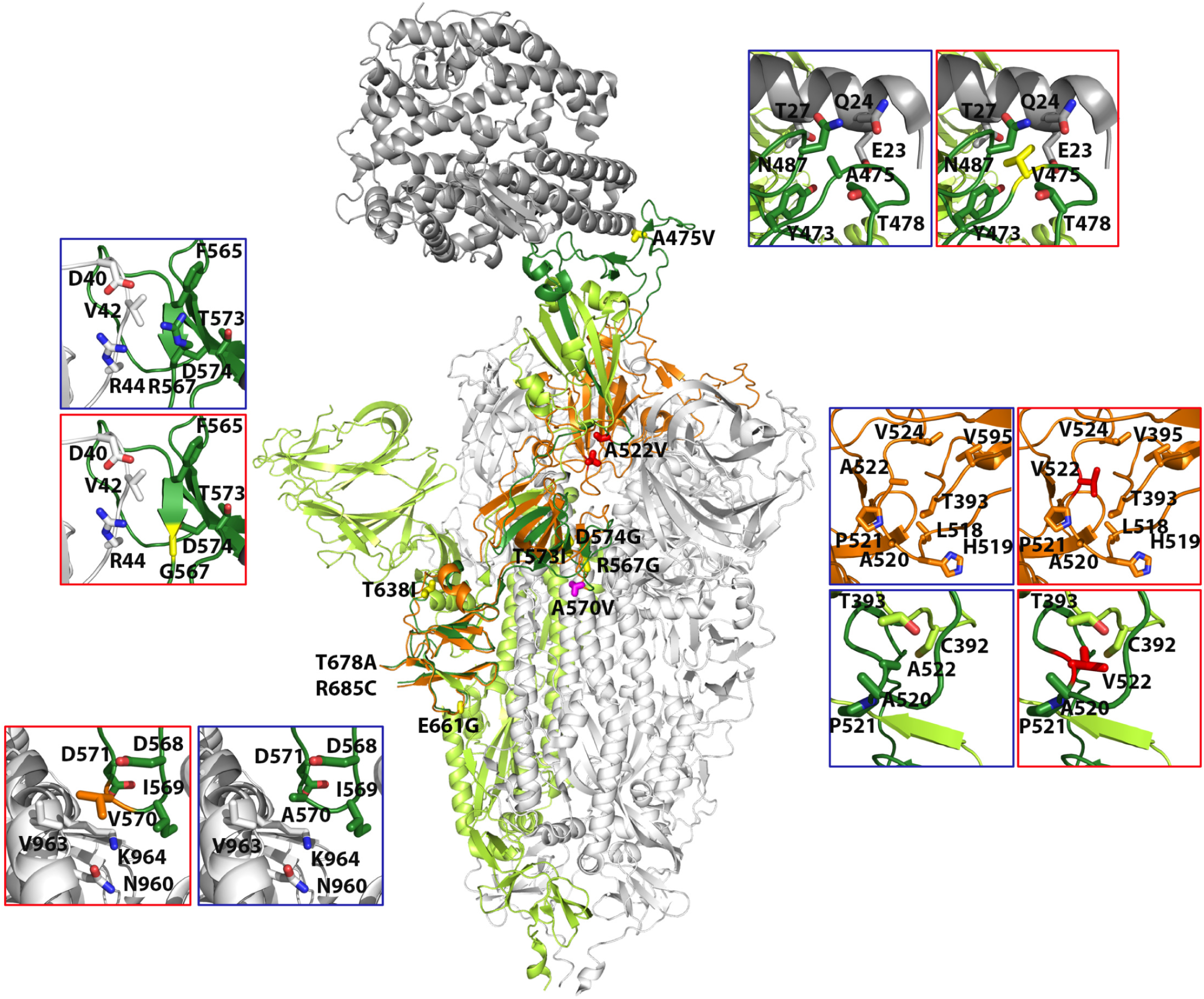
Location of amino acid substitutions in the three-dimensional structure of spike (S) protein. The central structure is a cartoon representation of S trimer (PDB code 7A94) with the reference monomer colored in green and dark-green, the latter marking the regions covered by amplicons A5-A6 (indicated in Fig. 1). The remaining monomers of the S trimer are shown in grey. The reference monomer contains de RBD domain in the “erect” position. A superimposition of this domain in the “open” conformation is also shown in orange. Substitutions are labeled, and amino acids are shown as sticks in different colors, according to associated disease category: exitus in red; moderate in magenta and mild in yellow. Insets highlight the interactions of some substituted positions with neighboring residues within a 5-Å radius. Except for position 522, two insets are shown per mutated position, indicating the original and mutated residues, squared in blue and red, respectively. For position 522, four insets are shown; the top two indicate the interactions of this residue in the open conformation of RBD, and the bottom two in the erect conformation. The substitutions, their frequency in the mutant spectrum, acceptability, functional score, and possible structural or functional effects are listed in Table 3.

### Deletion repertoire in SARS-CoV-2 mutant spectra

Deletions were also analyzed by UDS with a cut-off value of 0.5% (as detailed in Materials and Methods), with the same reads used for point mutations. The analyses identified five different deletions which spanned 3-13 nucleotides (nt) in the nsp12 (polymerase)-coding region, and five different deletions that spanned 2-51 nt in the S-coding region (Figs. 2 and S5 in https://saco.csic.es/index.php/s/8GH5aJgritCjEx5). In the nsp12 (polymerase)-coding region, the 4 nt and 13 nt deletions that disrupted the coding frame generated a stop codon 10 and 26 residues downstream, respectively. The 2 nt, 16 nt, 22 nt, and 28 nt deletions in the S-coding region led to stop codons 3 to 18 nucleotides downstream (Fig. S5 in https://saco.csic.es/index.php/s/8GH5aJgritCjEx5). The number of deletions that generated a stop codon was significantly higher in the S-coding region (26 out of 27 deletions) than in nsp12 (polymerase)-coding region (2 out of 10 deletions) (p<0.001; proportion test). The sites of deletions did not map in homopolymeric regions or tandem repeats, and they were not flanked by the same nucleotide types (Fig. S5 in https://saco.csic.es/index.php/s/8GH5aJgritCjEx5).

### Point mutation and deletion hot spots

The distribution of genomic variations (point mutations and deletions) per amplicon was similar for the four amplicons of the nsp12 (polymerase)-coding region (p-value > 0.05; proportion test). In contrast, amplicon A6 of the S-coding region accumulated higher number of total mutations than A5 (p<0.001; proportion test) (Fig. 7A). This difference may result from dissimilar functional constraints on the protein portions represented by each amplicon, i.e. a uniform distribution of polymerase motifs A to G among the four nsp12 (polymerase) amplicons, compared with the presence of the receptor-binding domain (RBD) in amplicon A5 of S (compare Figs. 1 and 7A).

**FIG 7.**
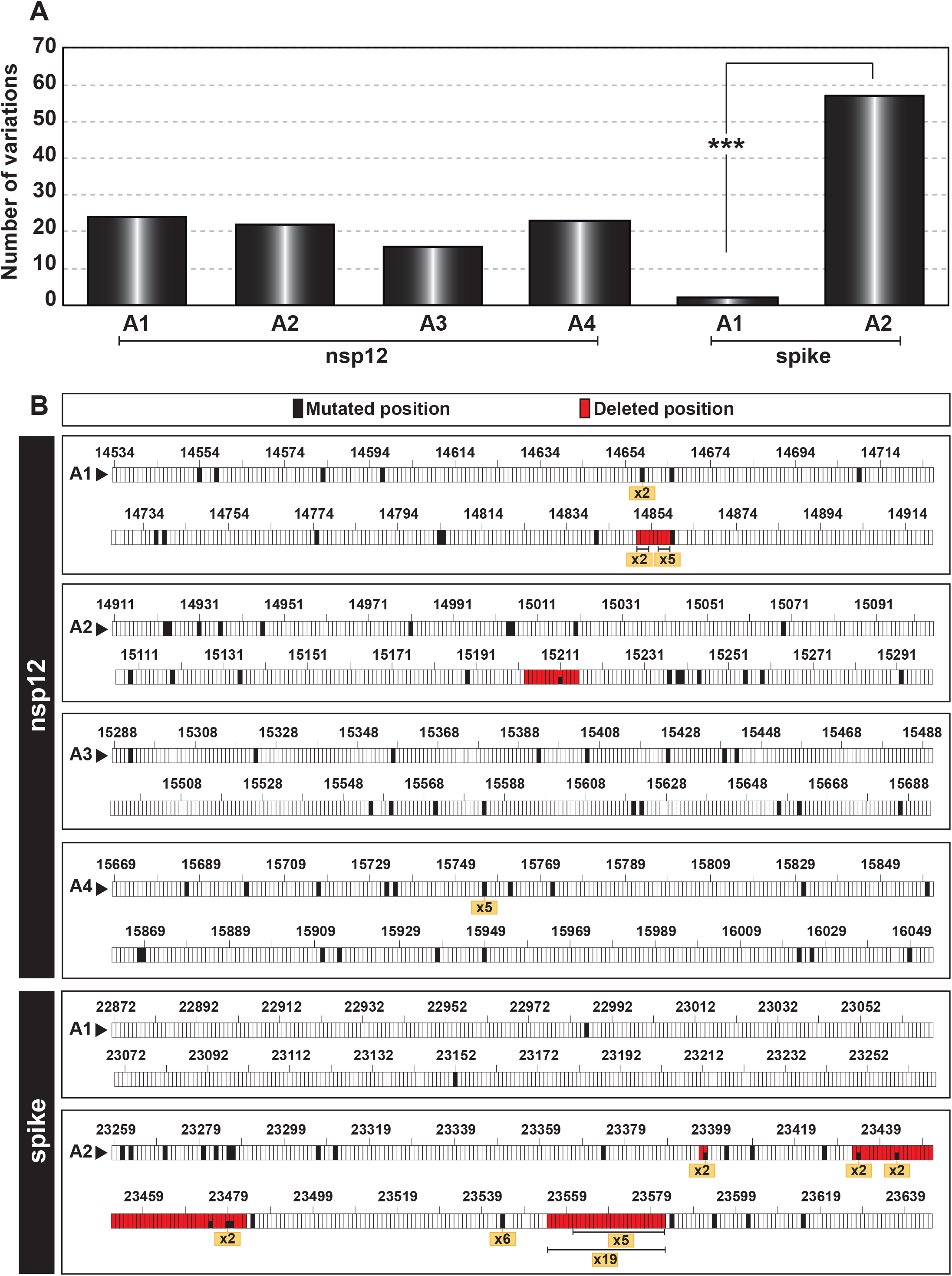
Point mutation and deletion hot spots in SARS-CoV-2 mutant spectra. **(A)** Distribution of the total number of different variations (point mutations and deletions, given in ordinate) among the amplicons analyzed (indicated in abscissa). The statistical significance of the differences was determined by the proportion test. ***, p<0.001; absence of connecting lines among nsp12 amplicons means that differences were not statistically significant. **(B)** Location of point mutations and deletions within each amplicon (indicated in each box). Genome residue numbering is according to reference NCBI accession number NC_045512.2. The numbers written in a yellow box refer to the number of patients whose virus carried the same mutation or deletion, and serve to identify hot spots. Point mutations and deletions were counted relative to the consensus sequence of the corresponding population.

Hot spots for SARS-CoV-2 variations have been described based on the comparison of consensus sequences of independent isolates (46–48). Here we have defined as hot spots those positions that presented the same point mutation or deletion in the mutant spectrum of at least five different isolates (Fig. 7B and Table S2 in https://saco.csic.es/index.php/s/8GH5aJgritCjEx5). Two hot spots were located in the nsp12 (polymerase)-coding region (a point mutation at position 15,756, and a deletion of residues 14,856 to 14,858), and two in the S-coding region (a point mutation at position 23,544 and a deletion of residues 23,555 to 23,582) (Fig. 7B). These hot spots do not coincide with those reported for SARS-CoV-2 consensus sequences (46–48).

### Geographical and temporal characterization of mutations based on CoV-GLUE database

SARS-CoV-2 mutant spectra from infected patients can include mutations that are also found as dominant in later isolates (27). In the mutant spectra of the 30 samples from our cohort, the ratio of amino acid substitutions (including those corresponding to divergence mutations) that were unique [not yet annotated in the CoV-GLUE database that is enabled by GISAID metadata (49)] versus those described in other (prior or subsequent) isolates was 0.2 (10 out of 60). Out of the 60 non-synonymous mutations, 8 (13.33%) were described worldwide at about the same time that they were identified in our cohort, and 19 (31.67%) were described afterwards (Fig. S6 in https://saco.csic.es/index.php/s/8GH5aJgritCjEx5). Of particular interest is S protein substitution S494P, located at the ACE-2 binding region (Table S2 in https://saco.csic.es/index.php/s/8GH5aJgritCjEx5) that reached epidemiological importance, and was found in some isolates of the alpha variant. Thus, SARS-CoV-2 mutant spectra ─in particular from patients that developed mild symptoms─ may constitute a rich reservoir of mutations with the potential to be represented in epidemiologically relevant variants.

## DISCUSSION

The UDS analysis of the nsp12 (polymerase)- and S-coding regions of 30 biological samples without cell culture passage confirm the presence of complex SARS-CoV-2 mutant spectra in diagnostic nasopharyngeal samples of the virus (23–28). Contrary to a previous conclusion with other patient cohorts (35, 36), our quantifications show that in both the nsp12 (polymerase)- and the S-coding regions analyzed there was a positive association between the number of point mutations and a mild disease manifestation in the corresponding patients. No such association was observed with the minority deletions that also populated the mutant spectra (Figs. 2 and 3). There are several non-mutually exclusive mechanisms that may contribute to a larger average number of point mutations in samples from patients that developed mild disease than in those from patients with moderate or severe disease. One is that the major sites of replication of the virus may not be identical in the three groups of patients. Mutational input may be affected by a variety of host cell functions, including editing activities (50), or as a consequence of the effects on polymerase fidelity of non-structural viral proteins that participate in genome replication, as evidenced with other RNA viruses (51–54). This possibility for SARS-CoV-2 is suggested by non-identical preferred transition mutation types in the isolates, depending on the associated disease severity (Table 1). A second influence may lie in a longer time of asymptomatic intra-host virus replication prior to the onset of mild symptoms and COVID-19 diagnosis. A prolonged replication time does not necessarily imply a larger viral load in the infected host, but it may entail an increase in the average number of variant genomes in the population. Another possibility is that bottleneck events ─which may transiently reduce the number of mutations scored within mutant spectra─ intervene with higher intensity in patients doomed to severe disease than those developing mild disease. This may come about through the immune response that may partially suppress viral replication, and that it is also part of the COVID-19 pathogenesis process (55–57). Several possibilities may explain dissimilar conclusions with other studies: (i) independent cohorts may have been infected by virus belonging to clades displaying non-identical behavior, and (ii) methodological differences such as in the criteria to classify patients according to COVID-19 symptoms, in the PCR-UDS resolution attained, or in the sample type taken for analysis (naso/oropharyngeal swabs versus nasopharyngeal aspirates), among others. The multiple factors that contribute to a mutant spectrum complexity beg for studies with other cohorts to try to clarify whether complexity of viral RNA in diagnostic samples responds to discernible virological parameters, and whether UDS data might help predicting disease evolution or response to treatment, as previously documented for hepatitis C (58, 59).

We have focused the mutant spectrum analysis on two regions of the SARS-CoV-2 genome whose encoded proteins are likely subjected to widely different constraints. The nsp12 (polymerase) is involved in genome replication and transcription, and the S glycoprotein has a major role in virus attachment, fusion and entry, as well as in defining the antigenic profile of the virus. A total of 41 different amino acid substitutions in nsp12 and 15 substitutions in S have been recorded in the 30 mutant spectra analyzed (Table S2 in https://saco.csic.es/index.php/s/8GH5aJgritCjEx5). Normalization to the sequenced protein length gives an average frequency of non-synonymous mutations of 8% for nsp12 and 6% for S in the mutant spectra. Three substitutions in S map in the receptor binding domain (RBD). One of them, A475V (present at 26% frequency in virus from a patient who developed mild disease) reduced the sensitivity to several monoclonal antibodies (60). S494P (dominant in virus from a patient who developed mild disease) was listed among the nine most frequent substitutions in a large-scale study of 506,768 SARS-CoV-2 isolates; it is considered a likely vaccine-escape substitution, and possibly involved also in increased transmissibility of some isolates of the alpha variant detected beginning September 2020 (61–63) (https://www.cdc.gov/).

Substitutions that are present at low frequency are associated with predicted more drastic structural and functional effects, and, some of them have been identified in the sequences compiled in the CoV-GLUE data base (compare Table 2 and Fig. S6 in https://saco.csic.es/index.php/s/8GH5aJgritCjEx5).

It is likely that disruptive amino acid substitutions belong to defective or minimally replicating (very low fitness) genomes that have either a transient existence in the population or that can be maintained at detectable levels by complementation (64) (for example those with lesions incompatible with polymerization activity). Defective genomes need not represent a biological or evolutionary dead end. They can exert modulatory effects on the entire population (65), and they also constitute a rich substrate for RNA recombination to rescue viable genomes that may become epidemiologically competent viruses.

Newly replicated genomes *in vivo* may incorporate deletions as a result of limited processivity of the coronavirus replicase (66, 67). Genomes with deletions may, on average, be subjected to stronger negative selection than genomes with point mutations, blurring differences in their frequency among samples from the three patient categories. This is likely to apply mainly to out of frame deletions that give rise to truncated proteins; for example, in the S-coding region we have identified deletion (Δ)23,555 to 23,570, Δ23,555 to 23,582, and Δ23,561 to 23,582 which are located near the S1/S2 cleavage site, and are expected to impair S function. Their maintenance to the point of reaching sufficient concentration to be detectable by UDS may reflect a higher efficiency of complementation of *trans*-acting structural proteins than non-structural proteins (64). This may also explain the lower frequency of out of frame deletions in the nsp12 (polymerase)- than in the S-coding region. It has been proposed that defective S proteins generated around the S1/S2 cleavage site could potentially reduce the severity of the infection (68).

All point mutations and deletions were found at frequencies below 30% in the corresponding mutant spectra. Several important biological and clinical features could influence the shape of SARS-CoV-2 mutant spectra. However, it should be considered that the large size of the coronavirus genome may limit the accumulation of mutations relative to less complex RNA genomes, due to negative effects of mutations on fitness (69). Not even the point mutation hot spots were found at frequencies above 1% in the quasispecies where they were present (compare Figs. 2 and 7). This is compatible with hot spots reflecting sites where lesions are more tolerated within a generally constrained RNA genome. The fact that hot spots according to mutant spectra do not coincide with those defined by consensus sequences adds to other observations that indicate that residue conservation criteria at these two levels do not coincide (70). That the great majority of mutations in SARS-CoV-2 mutant spectra are present at low frequency may slow down the response of the virus to specific selective constraints such as inhibitors or neutralizing antibodies. Under this scenario, viral load may become more important to furnish genomes with mutations required to respond to the constraints (71). Comparative measurements with different RNA viruses are needed to endorse these potential effects of mutant spectrum composition.

The higher percentage of transitions versus transversions, and of non-synonymous versus synonymous mutations is in agreement with previous reports of mutant spectrum and consensus sequence analyses of SARS-CoV-2 (23, 24, 35, 68, 72). Some differences with previous studies have been observed in the preferred mutation types (Fig. S3 in https://saco.csic.es/index.php/s/8GH5aJgritCjEx5). While in the mutant spectra of our cohort T to C was the most frequent point mutation, other studies reported C to T as the preferred mutation type (72, 73). C to T was, however, the most frequent mutation in virus from the subset of exitus patients (Fig. S3 in https://saco.csic.es/index.php/s/8GH5aJgritCjEx5) hinting at the possibility that in previous studies virus from patients with moderate and severe COVID-19 might have been over-represented. The lack of dominance of C to U transitions in our samples is also reflected in absence of depletion of amino acids A, H, Q, P and T when considering all amino acid substitutions observed (50, 74); the data of Fig. S3 in https://saco.csic.es/index.php/s/8GH5aJgritCjEx5 show a net gain of 3 amino acids in the A, H, Q, P and T subset. Another possible explanation for differences with previous studies could be that the latter focused on consensus sequences obtained from data bases covering the whole genome, whereas our results correspond to two specific genomic regions sequenced by UDS.

The six point mutations that altered the consensus sequence of the mutant spectra relative to reference NC_045512.2 (identified as “Divergence” in the heat map of Fig. 2 and in Table S2 in https://saco.csic.es/index.php/s/8GH5aJgritCjEx5) allowed an estimate of the rate of accumulation of mutations in the SARS-CoV-2 consensus sequence. The time interval between our Madrid isolates (dated April 2020) and the reference Wuhan isolate (dated December 2019) was 4 months. Considering this time interval, the average rate of evolution calculated is (1.6 ± 0.6) × 10^−3^ mutations per nucleotide and year (m/n/y), and it is only slightly higher than the average value from ten previous studies (1.2 ± 0.6) × 10^−3^ m/n/y (range 9.9 × 10^−4^ to 2.2 × 10^−3^ m/n/y) (73, 75–83). Higher evolutionary rates are frequently obtained the shorter is the time interval between the virus isolations considered for the calculation [reviewed in (84)]. The values for SARS-CoV-2 are comparable to those reported for other RNA viruses, suggesting that constraints at the quasispecies level may not affect significantly evolutionary rates considered at the epidemiological level (85). Our results hint at the possibility that SARS-CoV-2 evolving in patients exhibiting mild symptoms may contribute a majority of the variants that drive the high rates of evolution quantified at the epidemiological level.

## MATERIALS AND METHODS

### Patient cohort and stratification

Samples were collected during the first COVID-19 outbreak in Spain. The cohort of the study included 30 patients admitted to the Fundación Jiménez Díaz Hospital (FJD, Madrid, Spain) from April 3 to 29, 2020. All patients were confirmed to be positive for SARS-CoV-2 by a specific real-time RT-PCR (VIASURE Real Time PCR) with a Ct (cycle threshold, which is inversely correlated with viral RNA level) range of 15.6 to 28.5; the samples are a subset from the cohort that has been previously described in (37). Data collected included patient demographics, risk factors for COVID-19, and clinical information at the time of SARS-CoV-2 diagnosis (Table S1 in https://saco.csic.es/index.php/s/8GH5aJgritCjEx5). The parameters used to classify the patients included: (i) need of hospitalization, (ii) need of mechanical ventilation, (iii) admission to the intensive care unit (ICU), and (iv) exitus attributed to COVID-19. Taking these parameters into account, the patients were classified as mild, moderate and severe (exitus) cases according to the symptoms and hospitalization requirements: (i) mild symptoms (neither hospital admission nor ICU) (n=10), (ii) moderate symptoms (hospitalization without ICU) (n=10), and (iii) severe symptoms (hospitalization with admission to the ICU, and progression to exitus in all cases) (n=10). The clinical classification was established before the data analysis was performed.

### Oligonucleotide design

To design oligonucleotide primers, a total of 663 SARS-CoV-2 sequences from the NCBI database (https://www.ncbi.nlm.nih.gov/genbank/sars-cov-2-seqs/) were retrieved and aligned to the Wuhan-Hu-1 NCBI reference sequence NC_045512.2 (86). Nucleotide sequences were analyzed to design forward and reverse oligonucleotide primers (Table S4 in https://saco.csic.es/index.php/s/8GH5aJgritCjEx5). Four pairs of oligonucleotides were used for amplification and sequencing of four overlapping amplicons of the genomic region of nsp12 (polymerase) (nucleotides 14,511 to 16,075) encoding amino acids 366 to 871, and two pairs to cover the region of the S protein (nucleotides 22,853 to 23,666) encoding amino acids 438 to 694 (residue numbering according to reference sequence NC_045512.2) (Fig. 1 and Table S5 in https://saco.csic.es/index.php/s/8GH5aJgritCjEx5).

### RNA extraction and amplification of SARS-CoV-2 RNA from infected patients

SARS-CoV-2 RNA was extracted from 140 μl of medium from nasopharyngeal swabs using the QIAamp Viral RNA Mini Kit (250) (Qiagen), as specified by the manufacturer. Amplifications of nsp12 (polymerase)- and S-coding regions were performed by RT-PCR. Each region was amplified from 5 μl of the RNA preparation by RT-PCR using Transcriptor One Step RT-PCR kit (Roche Applied Science). To perform the RT-PCR, 5 μl of the preparation were mixed with 10 μl of 5x buffer, and 2 μl of a solution containing the forward primer, 2 μl of a solution with the reverse primer (50 ng/μl, each), and 1 µl of polymerase. Reaction parameters were 50°C for 30 min for the reverse transcription, an initial denaturing step at 94°C for 7 min, followed by 35 cycles of a denaturing step at 94°C for 10 s, an annealing step at 46-48°C for 30 s, an extension step at 68°C for 40 s, and then a final extension at 68°C for 7 min. In the case of samples with a Ct value greater than 26 (6 samples from the mild symptom group), the number of cycles was increased to 45. Negative controls (amplification reactions in the absence of RNA) were included in parallel to ascertain absence of contamination by template nucleic acids. Amplification products were analyzed by 2% agarose gel electrophoresis, using Gene Ruler 1 Kb Plus DNA ladder (Thermo Scientific) as molar mass standard. PCR products were purified (QIAquick Gel Extraction Kit, Qiagen), quantified (Qubit dsDNA Assay kit, Thermofisher Scientific), and tested for quality (TapeStation System, Agilent Technologies) prior to sequencing using the Illumina MiSeq platform. Dilutions of 1:10, 1:100 and 1:1,000 of the initial RNA preparation and subsequent amplification by RT-PCR were carried out for one patient of each disease severity (Fig. S7 in https://saco.csic.es/index.php/s/8GH5aJgritCjEx5). When amplification with the 1:1,000 dilution of template produced a visible DNA band, the ultra-deep sequencing analysis was performed with the undiluted template to avoid redundant copying of the same template molecules, as we have previously documented (87, 88).

### Ultra-Deep Sequencing of SARS-CoV-2 from Infected Patients

PCR products were adjusted to 4 × 10^9^ molecules/μl before generating DNA pools that were purified using Kapa Pure Beads (Kapabiosystems, Roche), and quantified using Qubit as previously described (38–40), and then fixed at 1.5 ng/μl. Purified DNA pools were further processed using the DNA library preparation kit Kapa Hyper Prep kit (Roche), during which each pool was indexed using SeqCap Adapter Kit A/B (Nimblegen) (24 Index). Each DNA pool was quantified by LightCycler 480, and sequenced using MiSeq sequencing platform with MiSeq Reagent kit v3 (2 × 300 bp mode with the 600 cycle kit) (Illumina).

### Bioinformatics analyses

Controls to stablish the basal error, the frequency of PCR-induced recombination, and the similarity of the results with different amplifications and sequencing runs were previously performed (38, 41, 89). Therefore, mutations identified with a frequency above the 0.5% cut-off value and with coverage greater than 10,000 reads were considered for the analyses, based on different controls carried out with hepatitis C virus (HCV), as detailed elsewhere (38, 90).

Beginning with the Fastq data, two bioinformatic pipelines [SeekDeep (42), and a new previously described pipeline for HCV (38)] were applied to HCV (Fig. S8 in https://saco.csic.es/index.php/s/8GH5aJgritCjEx5), and then adapted to SARS-CoV-2 to quantify deletions (termed VQS-Haplotyper, freely available in Github at this address https://github.com/biotechvana/VQS-haplotyper) (Fig. S9 in https://saco.csic.es/index.php/s/8GH5aJgritCjEx5). As control with an independent set of UDS data, we compared the point mutations and their frequencies within HCV quasispecies obtained using both bioinformatics procedures, and the results were very similar (r=0.9957 and p<0.0001; Pearson correlation test) (Fig. S8 in https://saco.csic.es/index.php/s/8GH5aJgritCjEx5). For SARS-CoV-2 mutant spectra, the analysis of clean reads using both pipelines yielded a robust similar number of point mutations and their frequencies (r=1 and p<0.0001; Pearson correlation test). Also, both pipelines produced similar results for deletions and their frequencies (r=0.4932 and p=0.0011; Pearson correlation test) (Fig. S9 in https://saco.csic.es/index.php/s/8GH5aJgritCjEx5). SeekDeep was applied using the following options: --extraExtractorCmds=-- checkRevComplementForPrimers – primerNumOfMismatches 3” “—extraProcessClusterCmds=--fracCutOff 0.005 – rescueExcludedOneOffLowFreqHaplotypes” (42). In the present study, point mutations, deletions and their frequencies were reported using SeekDeep, and diversity indices were calculated using VQS-Haplotyper followed by QSutils (43).

### Statistics

The correlation between results obtained by the bioinformatics pipelines was calculated using Pearson’s correlation. The statistical significance of difference between the number and type of mutations in mild, moderate and exitus patients as well as the differences between type of nucleotide changes and between PAM250 (accepted point mutations 250) and SNAP2 (Screening for Non-Acceptable Polymorphisms 2) values for amino acid substitutions were calculated by the proportion test. Statistics were inferred using software R version 4.0.2. The normality of data was tested with the Shapiro-Wilk normality test and the statistically significance of differences between diversity indices was calculated with a Wilcoxon test using GraphPad Prism 8.00.

### Data availability

The reference accession numbers of sequences retrieved from NCBI used to design oligonucleotide primers are given in Table S4 in https://saco.csic.es/index.php/s/8GH5aJgritCjEx5. Fastq files of SARS-CoV-2 samples included in the patient cohort are available in ENA under project id “PRJEB48766”. Nucleotide and amino acid replacements in SARS-CoV-2 from infected patients have been compiled in Table S2 in https://saco.csic.es/index.php/s/8GH5aJgritCjEx5.

## Ethics approval and consent to participate

This study was approved by the Ethics Committee and the Institutional Review Board of the FJD hospital (no. PIC-087-20-FJD).

## ACKNOWLEDGEMENTS

We acknowledge all personnel in the Clinical Microbiology Department of the FJD for help with the sample and data collection. We thank all health-care professionals who attended to COVID-19 patients, and collected the clinical samples that were included in this study in a difficult moment of the COVID-19 epidemic in Spain. We thank José María Aguado and Octavio Carretero for their support to the whole project. We are indebted to Cristina Villaverde for her technical expertise and help with the samples. We acknowledge Dres. J. Gregori and J. Quer for their contribution to the quasispecies analyses of HCV-infected samples.

This work was supported by Instituto de Salud Carlos III, Spanish Ministry of Science and Innovation (COVID-19 Research Call COV20/00181), and co‐financed by European Development Regional Fund ‘A way to achieve Europe’. The work was also supported by grants CSIC-COV19-014 from Consejo Superior de Investigaciones Científicas (CSIC), project 525/C/2021 from Fundació La Marató de TV3, PID2020-113888RB-I00 from Ministerio de Ciencia e Innovación, BFU2017-91384-EXP from Ministerio de Ciencia, Innovación y Universidades (MCIU), PI18/00210 and PI21/00139 from Instituto de Salud Carlos III and S2018/BAA-4370 (PLATESA2 from Comunidad de Madrid/FEDER). C.P., M.C. and P.M. are supported by the Miguel Servet programme of the Instituto de Salud Carlos III (CPII19/00001, CPII17/00006 and CP16/00116, respectively) cofinanced by the European Regional Development Fund (ERDF). CIBERehd (Centro de Investigación en Red de Enfermedades Hepáticas y Digestivas) is funded by Instituto de Salud Carlos III. Institutional grants from the Fundación Ramón Areces and Banco Santander to the CBMSO are also acknowledged. The team at CBMSO belongs to the Global Virus Network (GVN). B.M.-G. is supported by predoctoral contract PFIS FI19/00119 from Instituto de Salud Carlos III (Ministerio de Sanidad y Consumo) cofinanced by Fondo Social Europeo (FSE). R.L.-V. is supported by predoctoral contract PEJD-2019-PRE/BMD-16414 from Comunidad de Madrid. C.G.-C. is supported by predoctoral contract PRE2018-083422 from MCIU. BS was supported by a predoctoral research fellowship (Doctorados Industriales, DI-17-09134) from Spanish MINECO.

## REFERENCES

1. Huang C, Wang Y, Li X, Ren L, Zhao J, Hu Y, Zhang L, Fan G, Xu J, Gu X, Cheng Z, Yu T, Xia J, Wei Y, Wu W, Xie X, Yin W, Li H, Liu M, Xiao Y, Gao H, Guo L, Xie J, Wang G, Jiang R, Gao Z, Jin Q, Wang J, Cao B. 2020. Clinical features of patients infected with 2019 novel coronavirus in Wuhan, China. Lancet 395:497–506.

2. Dos Santos WG. 2021. Impact of virus genetic variability and host immunity for the success of COVID-19 vaccines. Biomed Pharmacother 136:111272.

3. Tillett RL, Sevinsky JR, Hartley PD, Kerwin H, Crawford N, Gorzalski A, Laverdure C, Verma SC, Rossetto CC, Jackson D, Farrell MJ, Van Hooser S, Pandori M. 2021. Genomic evidence for reinfection with SARS-CoV-2: a case study. Lancet Infect Dis 21:52–58.

4. To KK, Hung IF, Ip JD, Chu AW, Chan WM, Tam AR, Fong CH, Yuan S, Tsoi HW, Ng AC, Lee LL, Wan P, Tso E, To WK, Tsang D, Chan KH, Huang JD, Kok KH, Cheng VC, Yuen KY. 2021. COVID-19 re-infection by a phylogenetically distinct SARS-coronavirus-2 strain confirmed by whole genome sequencing. Clin Infect Dis 73(9):e2946–e2951.

5. Lee M. 2021. Lack of Severe Acute Respiratory Syndrome Coronavirus 2 Neutralization by Antibodies to Seasonal Coronaviruses: Making Sense of the Coronavirus Disease 2019 Pandemic. Clin Infect Dis 73:e1212–e1213.

6. Baum A, Fulton BO, Wloga E, Copin R, Pascal KE, Russo V, Giordano S, Lanza K, Negron N, Ni M, Wei Y, Atwal GS, Murphy AJ, Stahl N, Yancopoulos GD, Kyratsous CA. 2020. Antibody cocktail to SARS-CoV-2 spike protein prevents rapid mutational escape seen with individual antibodies. Science 369:1014–1018.

7. Hacisuleyman E, Hale C, Saito Y, Blachere NE, Bergh M, Conlon EG, Schaefer-Babajew DJ, DaSilva J, Muecksch F, Gaebler C, Lifton R, Nussenzweig MC, Hatziioannou T, Bieniasz PD, Darnell RB. 2021. Vaccine Breakthrough Infections with SARS-CoV-2 Variants. N Engl J Med 384:2212–2218.

8. McCallum M, Bassi J, De Marco A, Chen A, Walls AC, Di Iulio J, Tortorici MA, Navarro MJ, Silacci-Fregni C, Saliba C, Sprouse KR, Agostini M, Pinto D, Culap K, Bianchi S, Jaconi S, Cameroni E, Bowen JE, Tilles SW, Pizzuto MS, Guastalla SB, Bona G, Pellanda AF, Garzoni C, Van Voorhis WC, Rosen LE, Snell G, Telenti A, Virgin HW, Piccoli L, Corti D, Veesler D. 2021. SARS-CoV-2 immune evasion by the B.1.427/B.1.429 variant of concern. Science 373:648–654.

9. Nonaka CKV, Franco MM, Graf T, de Lorenzo Barcia CA, de Avila Mendonca RN, de Sousa KAF, Neiva LMC, Fosenca V, Mendes AVA, de Aguiar RS, Giovanetti M, de Freitas Souza BS. 2021. Genomic Evidence of SARS-CoV-2 Reinfection Involving E484K Spike Mutation, Brazil. Emerg Infect Dis 27:1522–1524.

10. Weisblum Y, Schmidt F, Zhang F, DaSilva J, Poston D, Lorenzi JC, Muecksch F, Rutkowska M, Hoffmann HH, Michailidis E, Gaebler C, Agudelo M, Cho A, Wang Z, Gazumyan A, Cipolla M, Luchsinger L, Hillyer CD, Caskey M, Robbiani DF, Rice CM, Nussenzweig MC, Hatziioannou T, Bieniasz PD. 2020. Escape from neutralizing antibodies by SARS-CoV-2 spike protein variants. Elife 9:e61312.

11. Domingo E, Perales C. 2019. Viral quasispecies. PLoS Genet 15:e1008271.

12. Domingo E, García-Crespo C, Perales C. 2021. Historical perspective on the discovery of the quasispecies concept. Annu Rev Virol 8:51–72.

13. Gregori J, Perales C, Rodriguez-Frias F, Esteban JI, Quer J, Domingo E. 2016. Viral quasispecies complexity measures. Virology 493:227–237.

14. Fuhrmann L, Jablonski KP, Beerenwinkel N. 2021. Quantitative measures of within-host viral genetic diversity. Curr Opin Virol 49:157–163.

15. Marcus PI, Rodriguez LL, Sekellick MJ. 1998. Interferon induction as a quasispecies marker of vesicular stomatitis virus populations. J Virol 72:542–549.

16. Farci P. 2001. Hepatitis C virus. The importance of viral heterogeneity. Clin Liver Dis 5:895–916.

17. Baranowski E, Ruiz-Jarabo CM, Pariente N, Verdaguer N, Domingo E. 2003. Evolution of cell recognition by viruses: a source of biological novelty with medical implications. Adv Virus Res 62:19–111.

18. Farci P. 2011. New insights into the HCV quasispecies and compartmentalization. Semin Liver Dis 31:356–74.

19. Domingo E, Sheldon J, Perales C. 2012. Viral quasispecies evolution. Microbiol Mol Biol Rev 76:159–216.

20. Honce R, Schultz-Cherry S. 2020. They are what you eat: Shaping of viral populations through nutrition and consequences for virulence. PLoS Pathog 16:e1008711.

21. Young DF, Wignall-Fleming EB, Busse DC, Pickin MJ, Hankinson J, Randall EM, Tavendale A, Davison AJ, Lamont D, Tregoning JS, Goodbourn S, Randall RE. 2019. The switch between acute and persistent paramyxovirus infection caused by single amino acid substitutions in the RNA polymerase P subunit. PLoS Pathog 15:e1007561.

22. Rima BK, Gatherer D, Young DF, Norsted H, Randall RE, Davison AJ. 2014. Stability of the parainfluenza virus 5 genome revealed by deep sequencing of strains isolated from different hosts and following passage in cell culture. J Virol 88:3826–36.

23. Karamitros T, Papadopoulou G, Bousali M, Mexias A, Tsiodras S, Mentis A. 2020. SARS-CoV-2 exhibits intra-host genomic plasticity and low-frequency polymorphic quasispecies. J Clin Virol 131:104585.

24. Jary A, Leducq V, Malet I, Marot S, Klement-Frutos E, Teyssou E, Soulie C, Abdi B, Wirden M, Pourcher V, Caumes E, Calvez V, Burrel S, Marcelin AG, Boutolleau D. 2020. Evolution of viral quasispecies during SARS-CoV-2 infection. Clin Microbiol Infect 26(11):1560.e1–1560.e4.

25. Rueca M, Bartolini B, Gruber CEM, Piralla A, Baldanti F, Giombini E, Messina F, Marchioni L, Ippolito G, Di Caro A, Capobianchi MR. 2020. Compartmentalized Replication of SARS-Cov-2 in Upper vs. Lower Respiratory Tract Assessed by Whole Genome Quasispecies Analysis. Microorganisms 8:E1302.

26. Capobianchi MR, Rueca M, Messina F, Giombini E, Carletti F, Colavita F, Castilletti C, Lalle E, Bordi L, Vairo F, Nicastri E, Ippolito G, Gruber CEM, Bartolini B. 2020. Molecular characterization of SARS-CoV-2 from the first case of COVID-19 in Italy. Clin Microbiol Infect 26:954–956.

27. Sun F, Wang X, Tan S, Dan Y, Lu Y, Zhang J, Xu J, Tan Z, Xiang X, Zhou Y, He W, Wan X, Zhang W, Chen Y, Tan W, Deng G. 2021. SARS-CoV-2 Quasispecies Provides an Advantage Mutation Pool for the Epidemic Variants. Microbiol Spectr 9:e0026121.

28. Andres C, Garcia-Cehic D, Gregori J, Pinana M, Rodriguez-Frias F, Guerrero-Murillo M, Esperalba J, Rando A, Goterris L, Codina MG, Quer S, Martin MC, Campins M, Ferrer R, Almirante B, Esteban JI, Pumarola T, Anton A, Quer J. 2020. Naturally occurring SARS-CoV-2 gene deletions close to the spike S1/S2 cleavage site in the viral quasispecies of COVID19 patients. Emerg Microbes Infect 9:1900–1911.

29. Wong YC, Lau SY, Wang To KK, Mok BWY, Li X, Wang P, Deng S, Woo KF, Du Z, Li C, Zhou J, Chan JFW, Yuen KY, Chen H, Chen Z. 2021. Natural Transmission of Bat-like Severe Acute Respiratory Syndrome Coronavirus 2 Without Proline-Arginine-Arginine-Alanine Variants in Coronavirus Disease 2019 Patients. Clin Infect Dis 73:e437–e444.

30. Xu D, Zhang Z, Wang FS. 2004. SARS-associated coronavirus quasispecies in individual patients. N Engl J Med 350:1366–7.

31. Tang JW, Cheung JL, Chu IM, Sung JJ, Peiris M, Chan PK. 2006. The large 386-nt deletion in SARS-associated coronavirus: evidence for quasispecies? J Infect Dis 194:808–13.

32. Liu J, Lim SL, Ruan Y, Ling AE, Ng LF, Drosten C, Liu ET, Stanton LW, Hibberd ML. 2005. SARS transmission pattern in Singapore reassessed by viral sequence variation analysis. PLoS Med 2:e43.

33. Park D, Huh HJ, Kim YJ, Son DS, Jeon HJ, Im EH, Kim JW, Lee NY, Kang ES, Kang CI, Chung DR, Ahn JH, Peck KR, Choi SS, Kim YJ, Ki CS, Park WY. 2016. Analysis of intrapatient heterogeneity uncovers the microevolution of Middle East respiratory syndrome coronavirus. Cold Spring Harb Mol Case Stud 2:a001214.

34. Borucki MK, Lao V, Hwang M, Gardner S, Adney D, Munster V, Bowen R, Allen JE. 2016. Middle East Respiratory Syndrome Coronavirus Intra-Host Populations Are Characterized by Numerous High Frequency Variants. PLoS One 11:e0146251.

35. Gregori J, Cortese MF, Pinana M, Campos C, Garcia-Cehic D, Andres C, Abril JF, Codina MG, Rando A, Esperalba J, Sulleiro E, Joseph J, Saubi N, Colomer-Castell S, Martin MC, Castillo C, Esteban JI, Pumarola T, Rodriguez-Frias F, Anton A, Quer J. 2021. Host-dependent editing of SARS-CoV-2 in COVID-19 patients. Emerg Microbes Infect 10:1777–1789.

36. Al Khatib HA, Benslimane FM, Elbashir IE, Coyle PV, Al Maslamani MA, Al-Khal A, Al Thani AA, Yassine HM. 2020. Within-Host Diversity of SARS-CoV-2 in COVID-19 Patients With Variable Disease Severities. Front Cell Infect Microbiol 10:575613.

37. Soria ME, Corton M, Martinez-Gonzalez B, Lobo-Vega R, Vazquez-Sirvent L, Lopez-Rodriguez R, Almoguera B, Mahillo I, Minguez P, Herrero A, Taracido JC, Macias-Valcayo A, Esteban J, Fernandez-Roblas R, Gadea I, Ruiz-Hornillos J, Ayuso C, Perales C. 2021. High SARS-CoV-2 viral load is associated with a worse clinical outcome of COVID-19 disease. Access Microbiol 3:000259.

38. Soria ME, Gregori J, Chen Q, Garcia-Cehic D, Llorens M, de Avila AI, Beach NM, Domingo E, Rodriguez-Frias F, Buti M, Esteban R, Esteban JI, Quer J, Perales C. 2018. Pipeline for specific subtype amplification and drug resistance detection in hepatitis C virus. BMC Infect Dis 18:446.

39. Soria ME, Garcia-Crespo C, Martinez-Gonzalez B, Vazquez-Sirvent L, Lobo-Vega R, de Avila AI, Gallego I, Chen Q, Garcia-Cehic D, Llorens-Revull M, Briones C, Gomez J, Ferrer-Orta C, Verdaguer N, Gregori J, Rodriguez-Frias F, Buti M, Esteban JI, Domingo E, Quer J, Perales C. 2020. Amino Acid Substitutions Associated with Treatment Failure for Hepatitis C Virus Infection. J Clin Microbiol 58:e01985–20.

40. Chen Q, Perales C, Soria ME, Garcia-Cehic D, Gregori J, Rodriguez-Frias F, Buti M, Crespo J, Calleja JL, Tabernero D, Vila M, Lazaro F, Rando-Segura A, Nieto-Aponte L, Llorens-Revull M, Cortese MF, Fernandez-Alonso I, Castellote J, Niubo J, Imaz A, Xiol X, Castells L, Riveiro-Barciela M, Llaneras J, Navarro J, Vargas-Blasco V, Augustin S, Conde I, Rubin A, Prieto M, Torras X, Margall N, Forns X, Marino Z, Lens S, Bonacci M, Perez-Del-Pulgar S, Londono MC, Garcia-Buey ML, Sanz-Cameno P, Morillas R, Martro E, Saludes V, Masnou-Ridaura H, Salmeron J, Quiles R, Carrion JA, Forne M, Rosinach M, Fernandez I, et al. 2020. Deep-sequencing reveals broad subtype-specific HCV resistance mutations associated with treatment failure. Antiviral Res 174:104694.

41. Perales C, Chen Q, Soria ME, Gregori J, Garcia-Cehic D, Nieto-Aponte L, Castells L, Imaz A, Llorens-Revull M, Domingo E, Buti M, Esteban JI, Rodriguez-Frias F, Quer J. 2018. Baseline hepatitis C virus resistance-associated substitutions present at frequencies lower than 15% may be clinically significant. Infect Drug Resist 11:2207–2210.

42. Hathaway NJ, Parobek CM, Juliano JJ, Bailey JA. 2018. SeekDeep: single-base resolution de novo clustering for amplicon deep sequencing. Nucleic Acids Res 46:e21.

43. Guerrero-Murillo M, Gregori i Font J. 2018. QSutils: Quasispecies Diversity. R package version 1.0.0.

44. Feng DF, Doolittle RF. 1996. Progressive alignment of amino acid sequences and construction of phylogenetic trees from them. Methods in Enzymol 266:368–82.

45. Hecht M, Bromberg Y, Rost B. 2015. Better prediction of functional effects for sequence variants. BMC Genomics 16 Suppl 8:S1.

46. Alouane T, Laamarti M, Essabbar A, Hakmi M, Bouricha EM, Chemao-Elfihri MW, Kartti S, Boumajdi N, Bendani H, Laamarti R, Ghrifi F, Allam L, Aanniz T, Ouadghiri M, El Hafidi N, El Jaoudi R, Benrahma H, Attar JE, Mentag R, Sbabou L, Nejjari C, Amzazi S, Belyamani L, Ibrahimi A. 2020. Genomic Diversity and Hotspot Mutations in 30,983 SARS-CoV-2 Genomes: Moving Toward a Universal Vaccine for the "Confined Virus"? Pathogens 9(10):829.

47. Badua C, Baldo KAT, Medina PMB. 2021. Genomic and proteomic mutation landscapes of SARS-CoV-2. J Med Virol 93:1702–1721.

48. Laamarti M, Alouane T, Kartti S, Chemao-Elfihri MW, Hakmi M, Essabbar A, Laamarti M, Hlali H, Bendani H, Boumajdi N, Benhrif O, Allam L, El Hafidi N, El Jaoudi R, Allali I, Marchoudi N, Fekkak J, Benrahma H, Nejjari C, Amzazi S, Belyamani L, Ibrahimi A. 2020. Large scale genomic analysis of 3067 SARS-CoV-2 genomes reveals a clonal geo-distribution and a rich genetic variations of hotspots mutations. PLoS One 15:e0240345.

49. Shu Y, McCauley J. 2017. GISAID: Global initiative on sharing all influenza data - from vision to reality. Euro Surveill 22:30494.

50. Mourier T, Sadykov M, Carr MJ, Gonzalez G, Hall WW, Pain A. 2021. Host-directed editing of the SARS-CoV-2 genome. Biochem Biophys Res Commun 538:35–39.

51. Smith EC, Case JB, Blanc H, Isakov O, Shomron N, Vignuzzi M, Denison MR. 2015. Mutations in coronavirus nonstructural protein 10 decrease virus replication fidelity. J Virol 89:6418–26.

52. Stapleford KA, Rozen-Gagnon K, Das PK, Saul S, Poirier EZ, Blanc H, Vidalain PO, Merits A, Vignuzzi M. 2015. Viral Polymerase-Helicase Complexes Regulate Replication Fidelity To Overcome Intracellular Nucleotide Depletion. J Virol 89:11233–44.

53. Agudo R, de la Higuera I, Arias A, Grande-Perez A, Domingo E. 2016. Involvement of a joker mutation in a polymerase-independent lethal mutagenesis escape mechanism. Virology 494:257–266.

54. Collins ND, Beck AS, Widen SG, Wood TG, Higgs S, Barrett ADT. 2018. Structural and Nonstructural Genes Contribute to the Genetic Diversity of RNA Viruses. mBio 9:e01871–18.

55. V’Kovski P, Kratzel A, Steiner S, Stalder H, Thiel V. 2021. Coronavirus biology and replication: implications for SARS-CoV-2. Nat Rev Microbiol 19:155–170.

56. Cheemarla NR, Watkins TA, Mihaylova VT, Wang B, Zhao D, Wang G, Landry ML, Foxman EF. 2021. Dynamic innate immune response determines susceptibility to SARS-CoV-2 infection and early replication kinetics. J Exp Med 218:e20210583.

57. Harrison AG, Lin T, Wang P. 2020. Mechanisms of SARS-CoV-2 Transmission and Pathogenesis. Trends Immunol 41:1100–1115.

58. Farci P, Shimoda A, Coiana A, Diaz G, Peddis G, Melpolder JC, Strazzera A, Chien DY, Munoz SJ, Balestrieri A, Purcell RH, Alter HJ. 2000. The outcome of acute hepatitis C predicted by the evolution of the viral quasispecies. Science 288:339–44.

59. Farci P, Strazzera R, Alter HJ, Farci S, Degioannis D, Coiana A, Peddis G, Usai F, Serra G, Chessa L, Diaz G, Balestrieri A, Purcell RH. 2002. Early changes in hepatitis C viral quasispecies during interferon therapy predict the therapeutic outcome. Proc Natl Acad Sci U S A 99:3081–6.

60. Li Q, Wu J, Nie J, Zhang L, Hao H, Liu S, Zhao C, Zhang Q, Liu H, Nie L, Qin H, Wang M, Lu Q, Li X, Sun Q, Liu J, Zhang L, Li X, Huang W, Wang Y. 2020. The Impact of Mutations in SARS-CoV-2 Spike on Viral Infectivity and Antigenicity. Cell 182:1284–1294 e9.

61. Grabowski F, Preibisch G, Gizinski S, Kochanczyk M, Lipniacki T. 2021. SARS-CoV-2 Variant of Concern 202012/01 Has about Twofold Replicative Advantage and Acquires Concerning Mutations. Viruses 13:392.

62. Alenquer M, Ferreira F, Lousa D, Valerio M, Medina-Lopes M, Bergman ML, Goncalves J, Demengeot J, Leite RB, Lilue J, Ning Z, Penha-Goncalves C, Soares H, Soares CM, Amorim MJ. 2021. Signatures in SARS-CoV-2 spike protein conferring escape to neutralizing antibodies. PLoS Pathog 17:e1009772.

63. Wang R, Chen J, Gao K, Wei GW. 2021. Vaccine-escape and fast-growing mutations in the United Kingdom, the United States, Singapore, Spain, India, and other COVID-19-devastated countries. Genomics 113:2158–2170.

64. Sola I, Almazan F, Zuniga S, Enjuanes L. 2015. Continuous and Discontinuous RNA Synthesis in Coronaviruses. Annu Rev Virol 2:265–88.

65. Vignuzzi M, Lopez CB. 2019. Defective viral genomes are key drivers of the virus-host interaction. Nat Microbiol 4:1075–1087.

66. Posthuma CC, Te Velthuis AJW, Snijder EJ. 2017. Nidovirus RNA polymerases: Complex enzymes handling exceptional RNA genomes. Virus Res 234:58–73.

67. Hillen HS, Kokic G, Farnung L, Dienemann C, Tegunov D, Cramer P. 2020. Structure of replicating SARS-CoV-2 polymerase. Nature 584:154–156.

68. Armero A, Berthet N, Avarre JC. 2021. Intra-Host Diversity of SARS-Cov-2 Should Not Be Neglected: Case of the State of Victoria, Australia. Viruses 13:133.

69. Domingo E, Schuster P. 2016. Quasispecies: from theory to experimental systems. Current Topics in Microbiology and Immunology. Vol. 392. Springer.

70. Garcia-Crespo C, Soria ME, Gallego I, Avila AI, Martinez-Gonzalez B, Vazquez-Sirvent L, Gomez J, Briones C, Gregori J, Quer J, Perales C, Domingo E. 2020. Dissimilar Conservation Pattern in Hepatitis C Virus Mutant Spectra, Consensus Sequences, and Data Banks. J Clin Med 9:3450.

71. Domingo E, Perales C. 2012. From quasispecies theory to viral quasispecies: how complexity has permeated virology. Math Model Nat Phenom 7:32–49.

72. Sarkar R, Mitra S, Chandra P, Saha P, Banerjee A, Dutta S, Chawla-Sarkar M. 2021. Comprehensive analysis of genomic diversity of SARS-CoV-2 in different geographic regions of India: an endeavour to classify Indian SARS-CoV-2 strains on the basis of co-existing mutations. Arch Virol 166:801–812.

73. Simmonds P. 2020. Rampant C-->U Hypermutation in the Genomes of SARS-CoV-2 and Other Coronaviruses: Causes and Consequences for Their Short- and Long-Term Evolutionary Trajectories. mSphere 5:e00408–20.

74. Danchin A, Marliere P. 2020. Cytosine drives evolution of SARS-CoV-2. Environ Microbiol 22:1977–1985.

75. Li X, Zai J, Zhao Q, Nie Q, Li Y, Foley BT, Chaillon A. 2020. Evolutionary history, potential intermediate animal host, and cross-species analyses of SARS-CoV-2. J Med Virol 92:602–611.

76. Nie Q, Li X, Chen W, Liu D, Chen Y, Li H, Li D, Tian M, Tan W, Zai J. 2020. Phylogenetic and phylodynamic analyses of SARS-CoV-2. Virus Res 287:198098.

77. Bai Y, Jiang D, Lon JR, Chen X, Hu M, Lin S, Chen Z, Wang X, Meng Y, Du H. 2020. Comprehensive evolution and molecular characteristics of a large number of SARS-CoV-2 genomes reveal its epidemic trends. Int J Infect Dis 100:164–173.

78. Lai A, Bergna A, Acciarri C, Galli M, Zehender G. 2020. Early phylogenetic estimate of the effective reproduction number of SARS-CoV-2. J Med Virol 92:675–679.

79. Nabil B, Sabrina B, Abdelhakim B. 2021. Transmission route and introduction of pandemic SARS-CoV-2 between China, Italy, and Spain. J Med Virol 93:564–568.

80. Pereson MJ, Mojsiejczuk L, Martinez AP, Flichman DM, Garcia GH, Di Lello FA. 2021. Phylogenetic analysis of SARS-CoV-2 in the first few months since its emergence. J Med Virol 93:1722–1731.

81. Castells M, Lopez-Tort F, Colina R, Cristina J. 2020. Evidence of increasing diversification of emerging Severe Acute Respiratory Syndrome Coronavirus 2 strains. J Med Virol 92:2165–2172.

82. Diez-Fuertes F, Iglesias-Caballero M, Garcia-Perez J, Monzon S, Jimenez P, Varona S, Cuesta I, Zaballos A, Jimenez M, Checa L, Pozo F, Perez-Olmeda M, Thomson MM, Alcami J, Casas I. 2021. A Founder Effect Led Early SARS-CoV-2 Transmission in Spain. J Virol 95:e01583–20.

83. Liu Q, Zhao S, Shi CM, Song S, Zhu S, Su Y, Zhao W, Li M, Bao Y, Xue Y, Chen H. 2020. Population Genetics of SARS-CoV-2: Disentangling Effects of Sampling Bias and Infection Clusters. Genomics Proteomics Bioinformatics 18:640–647.

84. Domingo E. 2020. Virus as Populations. Academic Press, Elsevier, Amsterdam. Second Edition.

85. Domingo E, Garcia-Crespo C, Lobo-Vega R, Perales C. 2021. Mutation Rates, Mutation Frequencies, and Proofreading-Repair Activities in RNA Virus Genetics. Viruses 13:1882.

86. Wu F, Zhao S, Yu B, Chen YM, Wang W, Song ZG, Hu Y, Tao ZW, Tian JH, Pei YY, Yuan ML, Zhang YL, Dai FH, Liu Y, Wang QM, Zheng JJ, Xu L, Holmes EC, Zhang YZ. 2020. A new coronavirus associated with human respiratory disease in China. Nature 579:265–269.

87. de Avila AI, Gallego I, Soria ME, Gregori J, Quer J, Esteban JI, Rice CM, Domingo E, Perales C. 2016. Lethal Mutagenesis of Hepatitis C Virus Induced by Favipiravir. PLoS ONE 11:e0164691.

88. Gallego I, Soria ME, Gregori J, de Avila AI, Garcia-Crespo C, Moreno E, Gadea I, Esteban J, Fernandez-Roblas R, Esteban JI, Gomez J, Quer J, Domingo E, Perales C. 2019. Synergistic lethal mutagenesis of hepatitis C virus. Antimicrob Agents Chemother 63:e01653–19.

89. García-Crespo C, Gallego I, Soria ME, De Ávila AI, Martínez-González B, Vázquez-Sirvent L, Lobo-Vega R, Moreno E, Gómez J, Briones C, Gregori J, Quer J, Domingo E, Perales C. 2021. Population disequilibrium as promoter of adaptive explora-tions in hepatitis C virus. Viruses 13:616.

90. Gregori J, Salicru M, Domingo E, Sanchez A, Esteban JI, Rodriguez-Frias F, Quer J. 2014. Inference with viral quasispecies diversity indices: clonal and NGS approaches. Bioinformatics 30:1104–1111.

